# Spatial and temporal characteristics of laboratory-induced *Anopheles coluzzii* swarms: shape, structure and flight kinematics

**DOI:** 10.1101/2024.03.25.586329

**Authors:** Bèwadéyir Serge Poda, Antoine Cribellier, Lionel Feugère, Mathurin Fatou, Charles Nignan, Domonbabele François de Sale Hien, Pie Müller, Olivier Gnankiné, Roch Kounbobr Dabiré, Abdoulaye Diabaté, Florian T. Muijres, Olivier Roux

## Abstract

*Anopheles* mosquitoes mate at sunset in aerial swarms. The development of mating-based methods for effective malaria vector control requires a good knowledge of the flight behaviour of *Anopheles* species in mating swarms. However, the process of how swarms are formed and maintained remains poorly understood. Here, we characterized the three-dimensional spatial and temporal flight kinematics of *Anopheles coluzzii* males swarming above a ground marker. We observed that the location, shape and volume of swarms were highly stereotypic, consistent over the swarming duration, regardless of the number of individuals in the swarm. The swarm had an elliptical cone shape, and we observed a differential spatial distribution of flight kinematics parameters within the swarm volume. Among these parameters, only swarm density varied with swarm size. Using a sensory system-informed model, we show that swarm location and shape can accurately be modelled based on visual perception of the marker. To control swarm height, swarming individuals maintain an optical angle of the marker ranging from 24° to 55°. Limiting the viewing angle deviation to 4.5% of the maximum value results in the observed elliptical cone swarm shape. We discuss the implications of these finding in mating success, speciation and for vector control.

## Introduction

Malaria mosquitoes are among the world’s deadliest animals. Each year, they cause millions of *Plasmodium*-infections in humans and hundreds of thousands of human deaths [1]. The primary malaria prevention method is to limit mosquito biting through vector control, which is mainly based on the use of insecticides through treated bed nets and indoor residual spraying [2,3]. These vector control methods are increasingly under pressure due to the ongoing rise in insecticide resistance in mosquitoes [4,5]. Consequently, further reduction of malaria cannot be achieved using insecticide-based prevention methods alone. Instead, in a context of integrated vector management, for effective malaria control, these conventional control methods need to be augmented with alternative complementary methods [6–8].

Alternative complementary methods based on mating biology are currently being developed for effective control of malaria and dengue in a broader context of vector-borne disease control. Promising candidates are sterile insect technique [9,10], genetically modified mosquitoes [11,12], or incompatible insect technique [13,14]. The effectiveness of these control techniques is strongly linked to the ability of the modified mosquitoes to mate successfully with wild individuals. Consequently, the development of these methods requires a good knowledge of the mating biology and ecology of target mosquitoes [15,16]. Unfortunately, we have currently limited knowledge of mosquito mating behaviour.

*Anopheles* mosquitoes are known to mate mainly in-flight in outdoor swarms. Mating swarms are formed by males at sunset, usually above a visual ground marker, and females fly into the swarm to find a mate [17–19]. However, the processes of how male mosquitoes form and maintain a swarm, as well as their flight behaviour in the swarm remain largely unknown [16,19–21]. This limited knowledge is partly due to the difficulty to study the behaviour of relatively small insects (∼5 mm), under outdoor low light conditions (∼3-5 lux), in a relatively short time period per day (∼20 min), and in large aggregations of individuals (up to thousands of males) in which individuals fly fast (about 0.2-2 m/s) [17–19,22,23]. As a result, previous studies based on direct observations have provided limited data on *Anopheles* swarming and mating behaviour.

Experimental approaches using machine-vision-based tracking systems in controlled laboratory or semi-field conditions can at least partially resolve these limitations, as such an approach can provide quantitative data on the movement kinematics of swarming mosquitoes in controlled conditions. The first videography-based studies focused on two-dimensional trajectories or three-dimensional positions of swarming mosquitoes in natural or laboratory-induced swarms [17,24–26]. Recent advancements in machine-vision with high spatial and temporal resolution videography, and automised tracking techniques have increased the degree of automation in data collection [27–29]. This allowed for multi-target tracking of the three-dimensional trajectories of swarming mosquitoes [23,30–32], and have made available large datasets for in-depth analyses [33,34].

These studies provided a quantitative description of several important traits of mating swarms that could be useful in assessing the ‘quality’ of mosquito strains in mosquito release-based methods of vector control [32], or raise new avenues in explaining the speciation process and the diversification in the *Anopheles gambiae* complex [23,26,30,31]. However, important questions are still unanswered, particularly with respect to the temporal and spatial dynamics of swarming, and how the swarming mosquitoes interact with environmental cues such as the visual ground marker and sunset cues.

Here, we assessed these questions by studying experimentally-induced male swarms of *Anopheles coluzzii* mosquitoes, a primary vector of malaria in West Africa. We used a custom-built videography-based tracking system that records the three-dimensional motion of swarming individuals. We determined how swarm kinematics changes throughout three swarming periods: at swarm formation, at the peak swarming activity, and at the end of swarming; and we quantified how this relates to the swarm marker and sunset cues. Finally, we developed a sensory-cue-inspired model that describes how swarm location, shape and size depend on visual input in the swarming male mosquitoes. The analysis of these data enabled us to formulate new hypotheses on swarming and mating behaviour in *Anopheles* mosquitoes.

## Methods

### Mosquitoes

Experiments were conducted with a newly established (F2) *Anopheles coluzzii* line. The mosquito line was established from wild females collected in Bama (11° 24′ 14″ N, 4° 24′ 42″ W), a village located 30 km north of Bobo-Dioulasso, Burkina Faso. Indoor-resting gravid females belonging to the *Anopheles* genus were collected using mouth aspirators and transferred to the insectarium. Females were placed individually in oviposition cups containing tap water. After oviposition, females were identified to species by routine PCR-RFLP [35]. Newly hatched larvae from females identified as *An. coluzzii* were pooled. Larvae were reared in tap water and fed *ad libitum* with Tetramin^®^ Baby Fish Food (Tetrawerke, Germany). Adult mosquitoes were held in 30 cm × 30 cm × 30 cm mesh-covered cages and provided *ad libitum* with a 5% glucose solution. Insectarium conditions were maintained at a temperature of 27±2 °C, a 70±10% relative humidity, and a 12 h:12 h light:dark cycle.

### Video tracking system

We used the Trackit 3D Fly technology platform (SciTrackS GmbH, Bertschikon, Switzerland [36]) to reconstruct the three-dimensional trajectories of the flying mosquitoes. Trackit 3D Fly is a machine-vision-based technology platform conceived to track multiple moving objects in three dimensions with real time monitoring [23,36–38]. The set-up consisted of:

i. a stereoscopic camera kit including two synchronized digital cameras (Basler acA2040-90um NIR, Germany), operating at 50 frames per second, at an image resolution of 4 megapixels and in the range of near-infrared (NIR) light frequency (CMV4000 CMOS sensor). Each camera was equipped with a Fujinon lens (Fujinon DV3.4×3.8SA-SA1, Japan) with which we controlled focus, zoom and aperture, and a NIR band-pass filter (Midwest Optical Systems BP850-25.4, USA) placed between the lens and camera sensor. The filter blocked light outside the NIR wavelength band. The two cameras were connected by a trigger cable to record frames synchronically.
ii. We used four Raytec lamps (Raytec pulsestar illuminators PSTR-I96-HV, United Kingdom) controlled by a controller (PSTR-PSU-4CHNL-HV, UK) to illuminate the tracking area with infrared (IR) light at a wavelength of 850 nm and a power of 880 W. The lights were set to maximal power in constant mode via the controller and the freeware GardasoftMaint (available from www.gardasoft.com).
iii. The videography tracking system was controlled using a desktop computer (Intel Core i7-8700 CPU @ 3.20 GHz, RAM 16.0 GB, Windows 10), to which the cameras and IR light controller were connected via USB 3.0. On the computer both the tracking software and camera calibration software (MATLAB R2017a, MathWorks Inc) were installed. The stereoscopic camera system was calibrated with the MATLAB Calibration Toolbox [39] using a checker-pattern board (Pixartprinting, Italy). During experiments, the cameras streamed video to the computer, from which the tracking software reconstructed the three-dimensional flight tracks using a Kalman filtering loop algorithm [40].

### Laboratory set-up

The laboratory set-up was located at the Institut de Recherche en Sciences de la Santé (IRSS), Bobo-Dioulasso, Burkina Faso (West Africa). The video tracking system was set up in a room designed to create optimal environmental conditions to trigger *Anopheles* swarming behaviour, as described in detail by Niang et al. [41]. In summary, the experimental room consisted of a white 5.1 m × 4.7 m × 3.0 m (length × width × height) windowless lab space equipped with a set of hardware for simulating sunset conditions. These included (i) a ceiling light dimmed from maximum light intensity to full darkness in 30 minutes, (ii) a 0.5 m high black horizon around the complete room perimeter, and (iii) a bright light directed at one of the 5.1 m walls mimicking sunset.

Swarms were produced in a 2.0 m × 0.7 m × 1.8 m (length × depth × height) transparent Plexiglas flight arena, located in the centre of the room (Figure 1*a,b*). On one side of the flight arena, we placed a 40 cm × 40 cm black cotton cloth on the floor, which we used as a visual swarm marker. The two cameras were mounted on 0.65 m high tripods and placed close to the sunset wall at 1.35 m from each other and 1.9 m from the flight arena wall. The stereoscopic cameras were placed such that flying mosquitoes could be tracked in three-dimensions across the complete flight arena. The four IR lamps were fixed on one of the 4.7 m lab walls, above each other (0 m, 0.7 m, 1.1 m and 1.6 m from the floor), at 1.35 m from the flight arena and illuminating the flight arena from the side at a wavelength of 850 nm and a power of 880 W. The desktop computer that controlled the cameras and IR lamps was located outside the room, allowing the researcher to assess and control the experiments without having to enter the experimental room.

**Figure 1:**
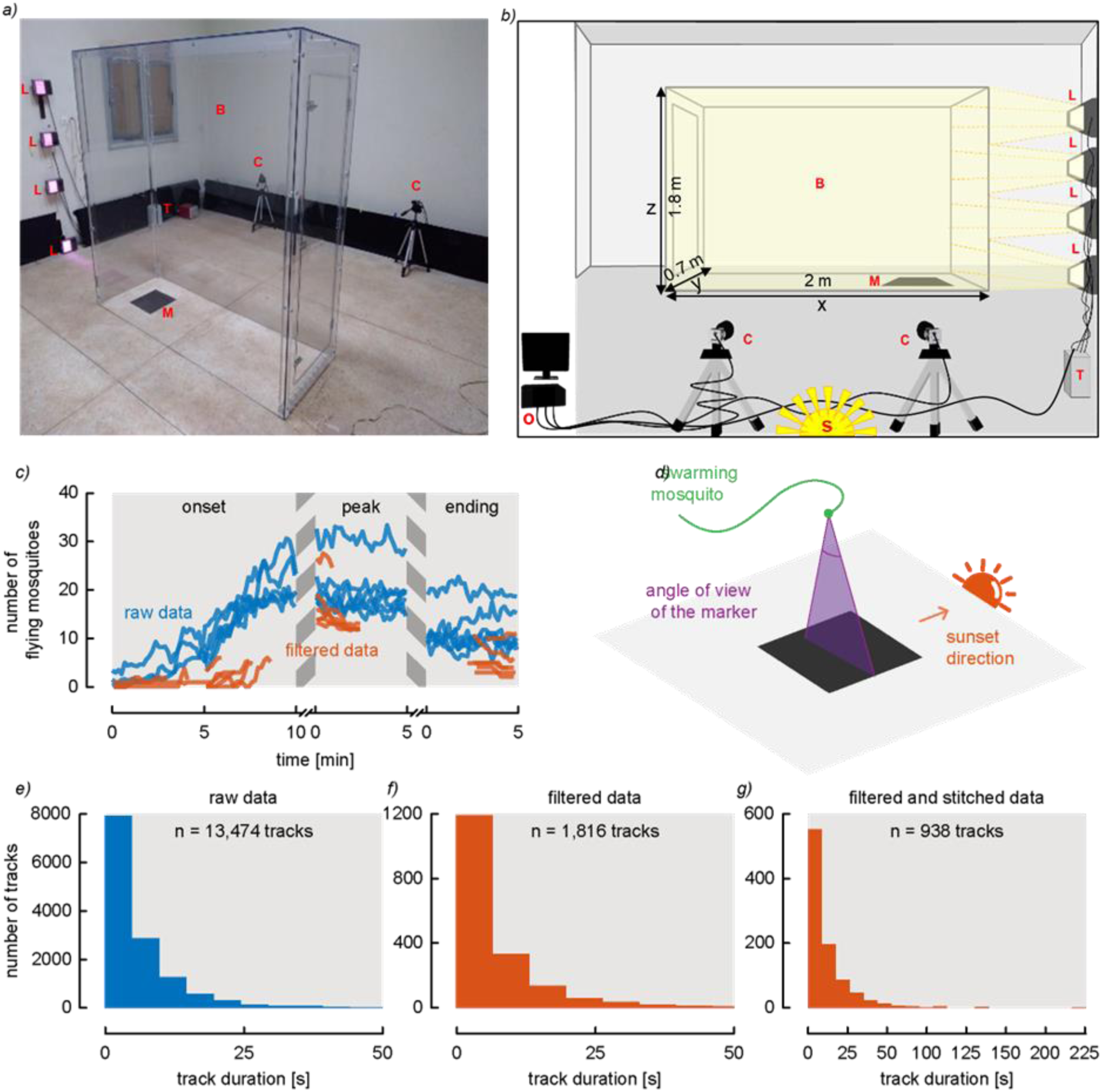
Experimental set-up for mosquito swarming behaviour studies. (*a,b*) Picture and schematic of the set-up, respectively. Capital letters highlight: (B) the Plexiglas box placed in the centre of the room; (M) a 40 cm × 40 cm black swarm marker; (C) two synchronised cameras located on (S) the sunset horizon side; (L) the four NIR lamps connected to (T) a controller. Cameras and lamp controller are connected to (O) a computer located outside the room. (*c*) Temporal dynamics of flight activity in the entire flight arena and in the swarms (in blue and red, respectively), during the three swarm phrases (start, peak and ending phases), for the six experimental replicates. (*d*) Schematic of our hypothesis that swarming mosquitoes use the optical angle of the ground marker and sunset direction to position themselves within the swarm. (*e-g*) Histograms of track duration of (*e*) all flight tracks, (*f*) selected swarming tracks and (*g*) stitched swarming trajectories of all the recordings.

**Figure 2:**
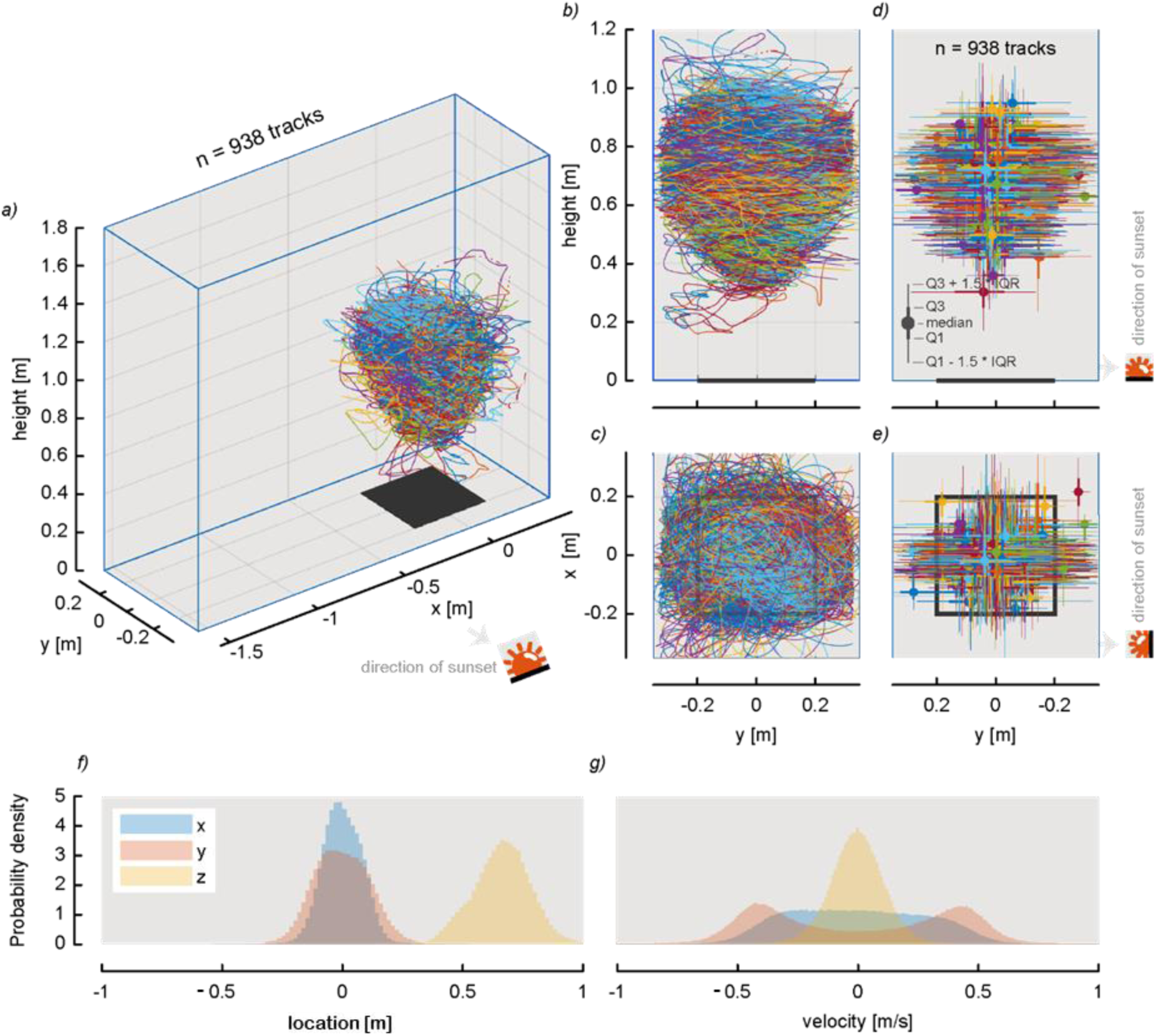
Swarming track locations and flight velocities. (*a-c*) All selected swarming tracks viewed (*a*) in three-dimensions, (*b*) from the side and (*c*) from the top (*n*_track_=938). (*d,e*) Side view (*d*) and top view (*e*) of the median location and quartiles of each swarming trajectory. The swarm marker is shown in black. (*f,g*) Distribution of locations (*f*) and velocities (*g*) of all swarming mosquitoes along the three axes (see legend in (*f*)).

### Experimental design for mosquito flight recording

For our swarming experiment, we used four-to six-day-old males of laboratory-reared *Anopheles coluzzii*. For each experiment, we released male mosquitoes into the flight arena one hour before swarming time, allowing them to acclimatize to the conditions. We chose to release 30 or 50 mosquitoes as this resulted in two swarm sizes of approximately 10 and 20 mosquitoes, respectively. Each swarming experiment was replicated three times (*i.e*. six experimental replicates in total).

At the start of each experiment (*i.e*. before the mosquitoes were released), both the ceiling and sunset lights were turned on. One hour after the mosquitoes were released, the ceiling lights dimmed slowly from full brightness to off during a period of 30 minutes. Meanwhile, the sunset light was kept at full power. Approximately 2-3 minutes after the ceiling lights went off, mosquitoes started to swarm, and the video tracking system was started.

During each experiment, we tracked the swarming mosquitoes at three key phases of the swarming period: (i) at the start of swarming, (ii) at peak swarming, and (iii) at the ending of the swarming activity (Figure 1*c*). We started the first recording when the first mosquito began a stationary looping flight over the swarm marker, and we continued this recording for 10 minutes. When the swarm reached its maximal size (∼5 minutes later), we started the second recording, which we continued for 5 minutes. To capture the decline in swarming activity, we started the third recording 5 minutes after the end of the second recording. This recording also continued for 5 minutes.

The tracking system was set to track up to 50 moving objects appearing as bright objects on a dark background, at a temporal resolution of 50 frames per second. The gain, detection threshold, exposure time and minimum object area were set at 5, 10, 10 ms and 20 pixels, respectively. During each data recording period, a video of the flying mosquitoes was recorded using Icecream Screen Recorder software (available from www.icecreamapps.com). These video recordings were used to check and validate the tracking accuracy (see the “Experimental data processing” section below).

### Experimental data processing

For each video recording, the tracking software provided a CSV file with the three-dimensional locations (*x,y,z*) of mosquitoes, at a temporal resolution of 20 ms. The coordinate system of the right-handed world reference frame has its origin (0,0,0) on the floor, at the centre of the ground marker. The *x*-axis is parallel to the sunset horizon, the *y*-axis points towards the sunset, and the *z*-axis vertically up. For convenience, we refer to the *z*-coordinates also as *height*. We defined global time *t* as the time relative to the start of swarming (*t*=0 s). Furthermore, we defined three time-metrics specific for the swarming phases (*t*_start_, *t*_peak_, *t*_end_, respectively), of which time is zero at the start of the phase-specific video recording (Figure 1*c*).

The output file contained the data of swarming and non-swarming flights of mosquitoes tracked throughout the flight arena. To select the data of interest (*i.e*. swarming flights), we plotted for each track the location along the three axes as a function of time, using the Trackit Data Selector program (SciTrackS Gmbh, Bertschikon, Switzerland) running on MATLAB (R2017a). On these graphs, we looked for swarming trajectories as sinusoidal tracks along the three axes (Figure S1*a,b*), as swarming behaviour has long been described in the literature as loop-like trajectories [17,22,24,30,31,42]. The swarming tracks were distinct from the non-swarming ones, and were manually selected (Figure S1*a,b*). To make sure we used data that are characteristic of the start, peak and end of swarming, and to avoid transition phases between these three phases, we selected the swarming flight tracks in the first 450 seconds, first 150 seconds and last 150 seconds of the recordings of the swarm start, peak and ending, respectively (see Figure 1*c*).

One of the main limitations of video tracking data of mosquito swarms is the interruptions of flight trajectories [22,30]. This results in a significant loss of information of the individual movements in the swarms, and is therefore a limitation in data analyses [22]. For these reasons, we used a specially designed application in Shiny R (*version* 4.3.1) (RMosquito, by SPILEn, France) to automatically reconstruct the interrupted trajectories, wherever possible (see Figure S1*c,d*). The RMosquito program used the principle of clustering to combine segments of interrupted trajectories. According to the three-dimensional spatial and temporal information, a score was established for each track. Then, a hierarchical clustering algorithm grouped the tracks into several clusters according to their score. Finally, the clusters were refined using four user parameters, including (i) minimal track duration, (ii) maximal distance to the nearest track point, (iii) maximal time difference with the nearest track point, and (iv) minimal duration of combined tracks. The reconstructed trajectories were visually checked for combination errors using three-dimensional plots and the video recordings, and the user parameter values were adjusted for correction. After testing several combinations of user parameter values, the following values were used to reconstruct the interrupted swarming trajectories: 0 s, 0.1 m, 0.2 s and 0 s for the minimal track duration, maximal distance to the nearest track point, maximal time difference with the nearest track point and minimal duration of combined tracks, respectively.

For the six experimental replicates, we reconstructed in total 13,474 flight tracks (Figure 1*e*). These tracks were from both flying non-swarming mosquitoes and mosquitoes that exhibited distinct swarming behaviours above the ground marker. Manual selection resulted in the identification of 1,816 swarming tracks, and consecutive automatic stitching of these swarming tracks resulted in a final dataset of 938 swarming flight trajectories (Figures 1*f,g* and S1*e,f*). The duration of these swarming flight tracks varied from a couple of seconds for the short tracks, to a maximum of 233 seconds for the longest track. Of these 938 swarming flight trajectories, 60 occurred during the swarm onset phase, 705 during the swarm peak phase, and 173 during swarm ending. This dataset was analysed to characterize the temporal dynamic of the flight kinematics of individual swarming mosquitoes and, characterize and visualize the three-dimensional shape and structure of the swarms.

### Characterising the flight kinematics of individual swarming mosquitoes

The selected and reconstructed tracks were then analysed in MATLAB R2022b. First, the three-dimensional trajectories were smoothed using a Savitzky-Golay filter [43], resulting in the smoothed flight trajectory array of positional vectors ([*x*(*t*), *y*(*t*), *z*(*t*)]). We then determined the temporal dynamics of velocity ([*u*(*t*), *v*(*t*), *w*(*t*)]) and acceleration ([*a*_*x*_(*t*), *a*_*y*_(*t*), *a*_*z*_(*t*)]) by taking the first and second temporal derivative of position, respectively. For this, we used a central finite difference scheme of second-order accuracy [44]. From the velocity data, we estimated the flight speed as 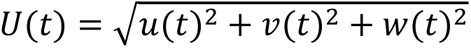, and from the acceleration vectors we estimated the corresponding net acceleration magnitude as 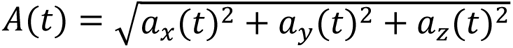 throughout each flight track. Furthermore, we estimated for each swarming mosquito the distance to closest neighbour at each instant of its swarming flight (*d*_neighbour_(*t*)).

From these kinematics time series, we determined for each swarming mosquito the average and standard deviation of all swarming kinematics parameters. These include per swarming mosquito, the average and standard deviation of location ((*x̅*, *y̅*, *z̅*), *σ*(*x*, *y*, *z*)), flight speed (*U̅*, *σ*(*U*)), acceleration (*A̅*, *σ*(*A*)), and the average of the distance to closest neighbour (*d̅*_neighbour_).

### Characterising and visualising the kinematics of the complete swarm

Next to the kinematics of individual swarming mosquitoes, we also quantified the kinematics of the swarm as a whole, to characterise the emergent swarming dynamics that results from the individual flight behaviours. Here, we defined the three-dimensional location and spread of the swarm as the average and standard deviation of the positional data of all mosquitoes in the swarm as (*x*_swarm_, *y*_swarm_, *z*_swarm_) and *σ*(*x*_swarm_, *y*_swarm_, *z*_swarm_), respectively.

Furthermore, we quantified the three-dimensional size and shape of the swarms as the volume containing 95% of the detected flight track positions. The remaining 5% of mosquito positions were considered as outliers (*i.e.* outside the swarm). To do so, we divided the two-dimensional surface of each view (*i.e*. front, side and top views) into an array of 2 cm × 2 cm cells. Then, we computed the depths that contained 95% of data inside each of these cells (Figure 3*a-c*). For this, we estimated the 2.5% and 97.5% quantiles of the third dimension normal to the view of interest (*i.e*. the *z* dimension when looking at the top view), and we defined the 95% depth as the distance between these two quantiles. Multiplying these depths by the area of cells allowed us to estimate the volume of the swarm that contain 95% of the swarming tracks (Figure 3*a*-*c*).

**Figure 3:**
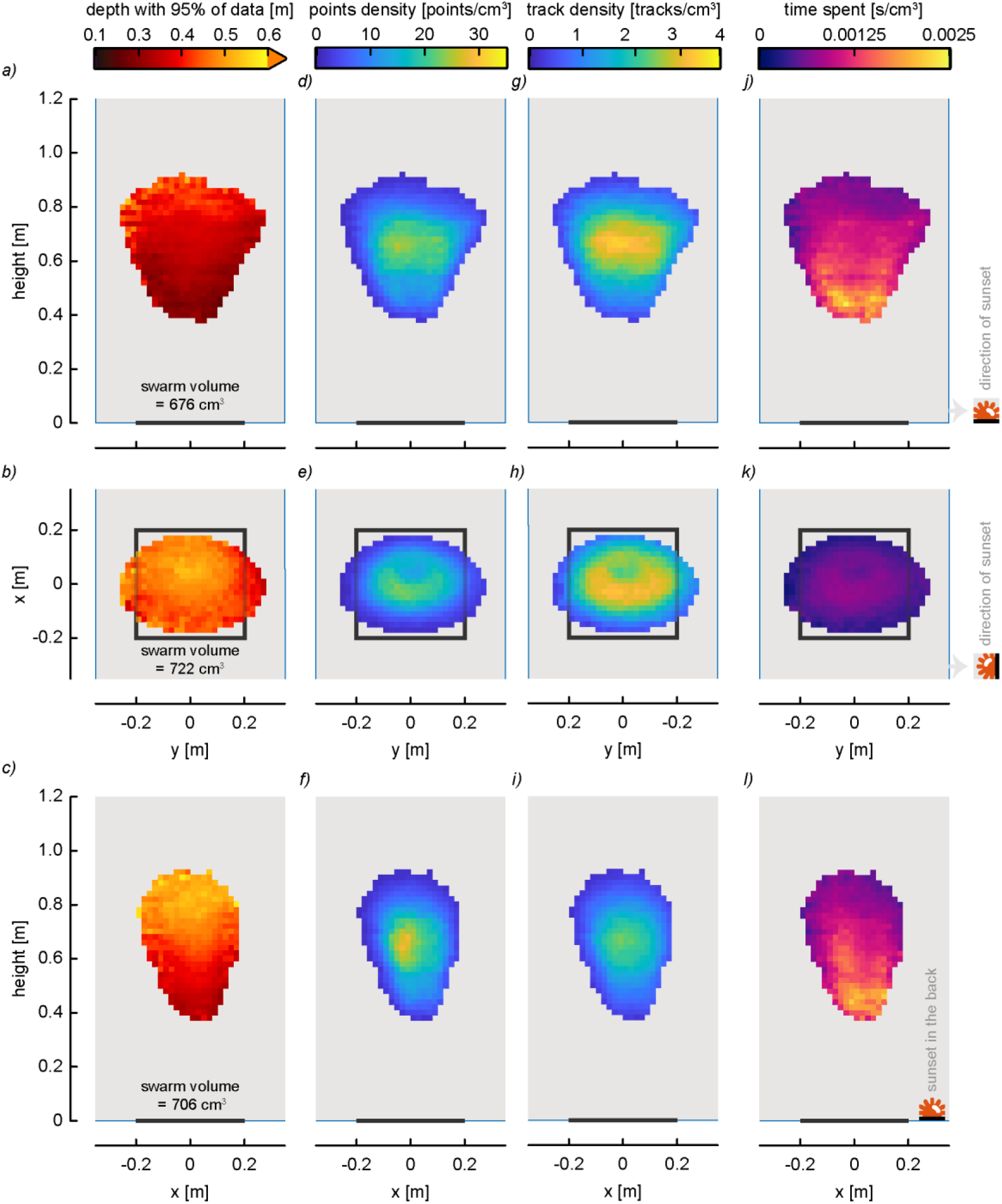
Swarm shape and structure. (*a-c*) Spatial distributions of the depth of each sub-volume comprising of 95% of the three-dimensional tracking data. These heatmaps describe the three-dimensional shape of the swarms. (*d-i*) Spatial distributions of the point density (*d-f*) and the track density (*g-i*) per cm^3^. (*j-l*) Spatial distribution of the mean time spent in each cm^3^ within the swarm. Cell with less than an average of one track per swarm and per recording (*i.e*. 6 swarms × 3 recordings = 18) were filtered out. The top, middle and bottom row show the swarm viewed from parallel to the sunset horizon, the top and normal to the sunset horizon, respectively (see sunset symbol). The swarm marker is shown as a black line or square.

Within the identified swarm volume, we visualised various swarming metrics using a set of three heatmaps, showing the swarm from the front, side and top (Figures 3, 4, S3 and S4). To do so, we divided the experimental volume into smaller rectangular sub-volumes of 2 cm × 2 cm × *depth* (Figure S2*a*), and within it we computed metrics such as the number of tracks in each of these sub-volumes (Figure S2*b*). Finally, we projected the results of each metric into two-dimensional heatmaps (Figure S2*c*). We did so for the three orthogonal planes (front, side and top views) to visualise the three-dimensional aspect of mosquito swarming behaviour (Figures 3, 4, S3 and S4). The different metrics computed are point and track densities (*i.e*. number of points or tracks per sub-volume), average time spent in each sub-volume, mean distance to closest neighbour, mean flight speed, and mean acceleration.

**Figure 4:**
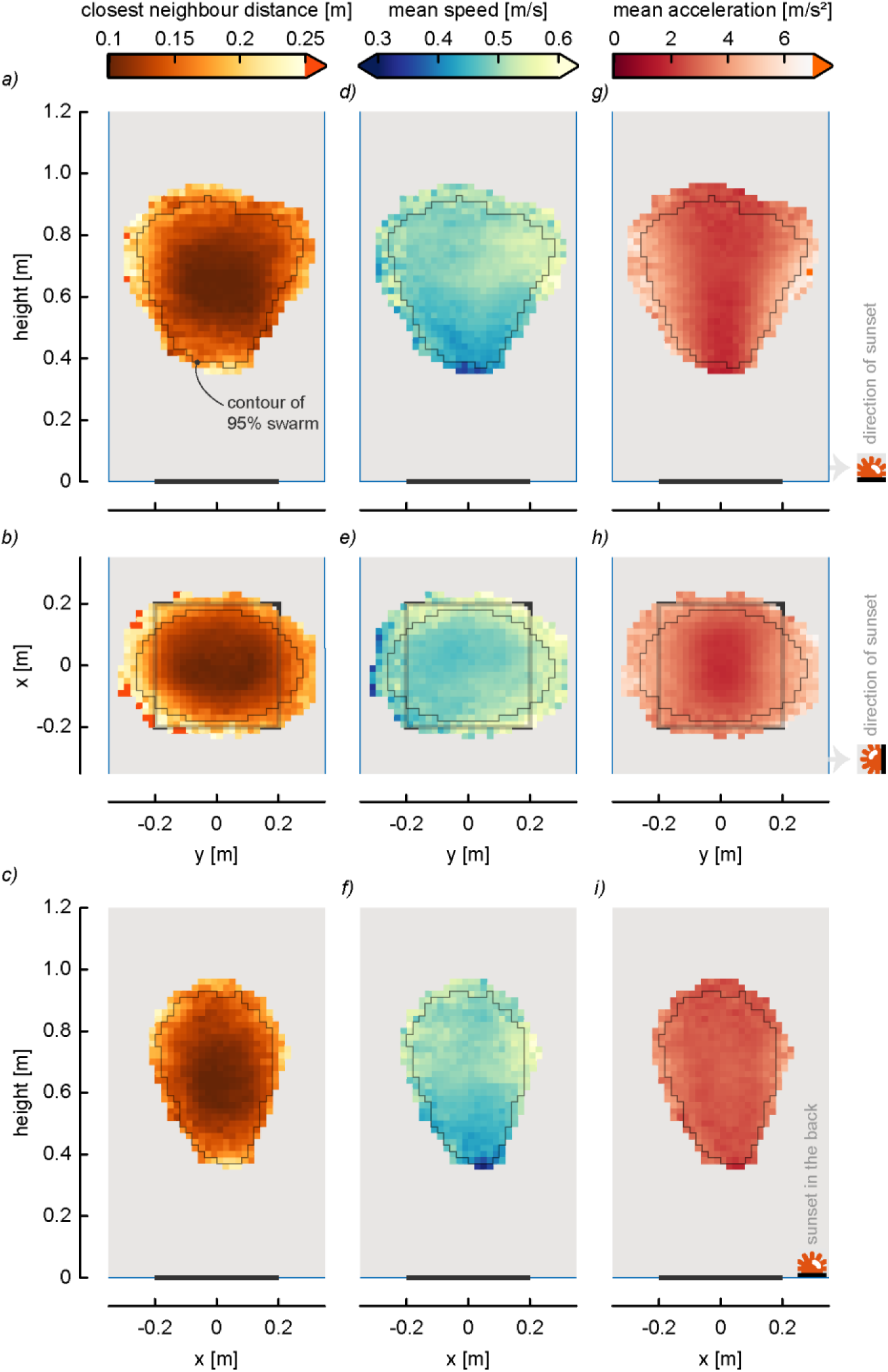
Spatial distribution of swarming flight kinematics. (*a-c*) Spatial distribution of the mean distance to the closest neighbour. (*d-i*) Spatial distribution of the mean flight speed (*d-f*) and mean acceleration (*g-i*), for mosquitoes swarming in each 2 cm × 2 cm cell. The top, middle and bottom row show the swarm viewed from parallel to the sunset horizon, the top and normal to the sunset horizon respectively (see sunset symbol). The swarm marker is shown as a black line or square.

The swarms are found to have quite variable depths depending on the chosen views. Therefore, it is unfair to compare point or track densities in one view to another view (*e.g*. 60 tracks in a 50 cm deep sub-volume *vs*. 30 tracks in a 25 cm deep sub-volume). Thus, to better compare metrics between the different two-dimensional views, we normalized point density, track density and average time spent by computing these metrics per cm^3^ in each sub-volume that contains 95% of the data (Figure 3*d-l*).

### Modelling mosquito swarming behaviour

In this study, we hypothesised that mosquitoes use visual cues of both the ground marker and sunset horizon to initiate swarming and maintain their position in the swarm. The ground marker can be used by swarming mosquitoes to control their horizontal position (*x,y*) and height (*z*) during swarming, and the sunset location allows mosquitoes to orient the main axis of the swarm.

The visual system of insects can accurately decode the optical angular size of objects as an estimate of the apparent size of the objects [45,46]. The angular size of objects scales with the ratio of object size and distance, and so flying insects use optical angular size and its temporal derivative (optical expansion) to control many aspects of flight, including navigation, landing manoeuvres, collision avoidance and evasive manoeuvres [47–49].

Here, we hypothesize that swarming mosquitoes use the optical angle of the ground marker as a metric for positioning themselves above the marker, both in height and horizontal location. Moreover, we suggest that mosquitoes also use the retinal position of the sunset to select the angle of view of the marker used to orient the main axis of the swarm (Figure 1*d*).

To test this hypothesis, we developed a visual-cue-inspired model of mosquito swarming, and compared the model output in the form of swarm position, shape and size with the experimental results. The model uses three behavioural rules to define the boundaries of the mosquito swarm (Figure 5*a*):

1. The optical angle of the ground marker decreases with increasing flight height, and so for our model we assume that the top and bottom boundaries of the swarm are set by minimal and maximal optical marker angle thresholds perceived by the swarming mosquitoes, respectively.
2. When a mosquito flies horizontally from a position right above the marker towards the edge of the marker, the optical angle of the marker decreases, thus causing a so-called optic contraction. We suggest that a swarming mosquito uses these optic contraction cues to trigger a turn, causing it to remain above the marker.
3. The swarming mosquitoes use the optical position of the sunset to orient themselves relative to the sunset, allowing them to discriminate between optical marker angles in the different orientations (parallel and normal to the sunset).

**Figure 5:**
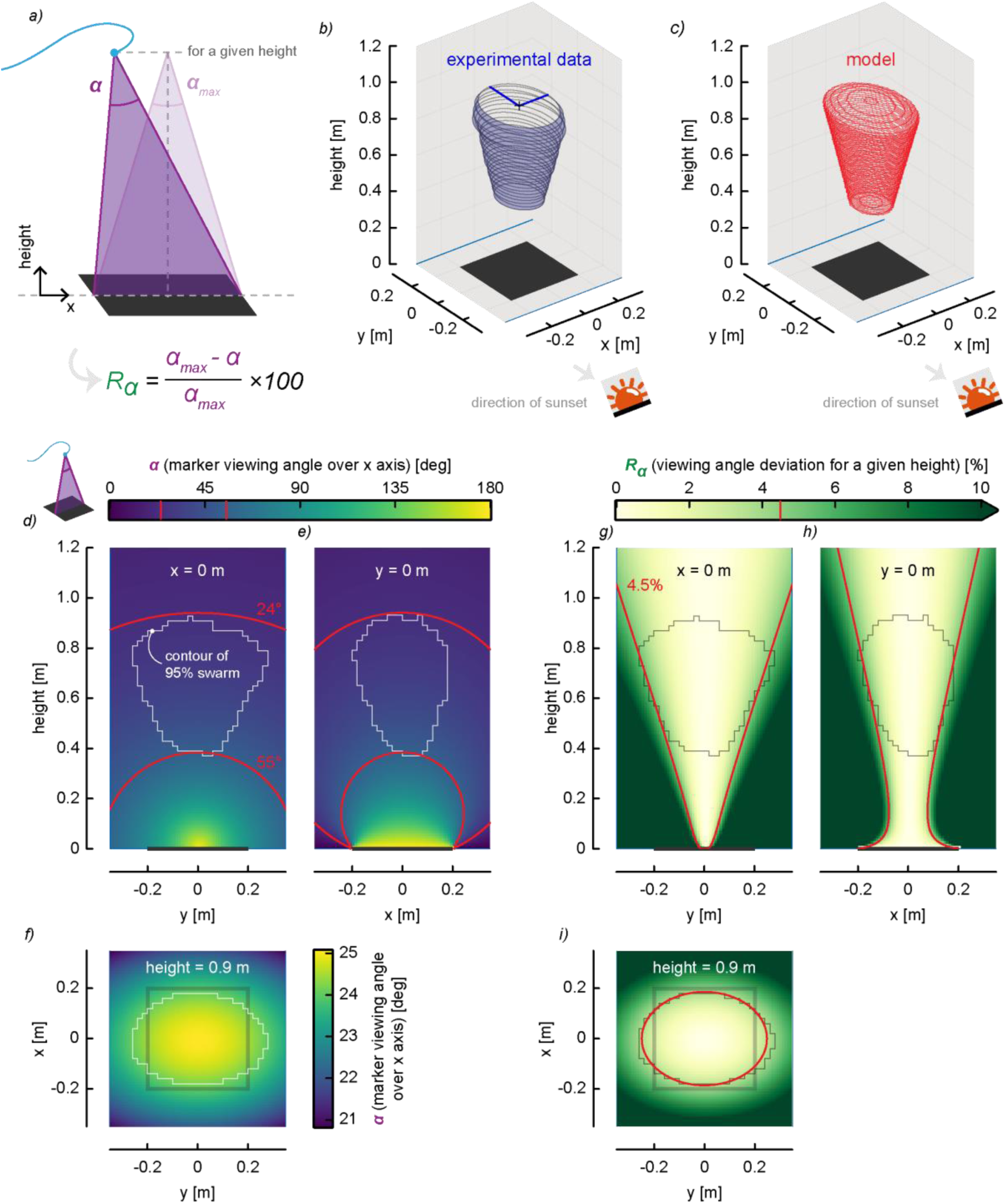
Modelling the marker viewing angle and the three-dimensional swarm shape. (*a*) Schematic showing how the viewing angle of the marker *α* and the viewing angle deviation *R*_α_ are defined at a given height. (*b,c*) The three-dimensional shape of the swarm based on experimental data (*b*), and on our mosquito vision-informed model (*c*). Both consist of a set of ellipses, based on (*b*) the experimental swarm radiuses along the *x*-axis and *y*-axis, and (*c*) on the swarming model parameters shown in (*d-i*). (*d-f*) The marker viewing angle along the *x*-axis (*α*) throughout 3D space, viewed from parallel to the sunset horizon, normal to the sunset horizon and top, respectively. The red and white lines show viewing angle iso-lines (*α*=24° and *α*=56°) and the swarm contour, respectively. (*g-i*) The percentage difference *R*_α_ between the maximum viewing angle over *x*-axis and current viewing angle for a given height, viewed from parallel to the sunset horizon, normal to the sunset horizon and top, respectively. The red and grey lines show iso-lines of *R*_α_=4,5% and the swarm contour, respectively. The swarm marker is shown as a black line or square.

To optimize the swarming model and fit it to the experimental data, we computed the optical marker angles throughout the swarming area and estimated the threshold angles for the model. These are the minimum and maximum optical angle (rule 1), and the reduction in optical angle relative to the maximum at a given height (rule 2). We then assessed the model accuracy by comparing the shapes of the model swarm with the experimental swarm shape.

### Statistical analyses

All statistical analyses were performed using R (*version* 4.3.1). We used Linear Mixed-Effects Models (*lmer* function, *lme4* package in R) to test how various motion parameters differ between the three-dimensional axes (*x,y,z*), the swarming activity phases (start, peak and end of swarming), and the swarm sizes at swarming peak. Furthermore, we analysed the interaction between three-dimensional axes and swarming activity phases (axes × swarming phases) to test whether the *x*, *y* and *z* components of motion parameters varied over the three swarming phases. The motion parameters that we tested were (i) the mean three-dimensional location of the swarming mosquitoes (*x̅*, *y̅*, *z̅*), (ii) the standard deviation of the swarming mosquito location *σ*(*x*, *y*, *z*), (iii) the mean flight speed of the swarming mosquitoes *U̅*, (iv) their mean acceleration magnitude *A̅*, and (v) the mean distance to closest neighbours of the swarming mosquitoes *d̅*_neighbour_. The experimental swarm numbers (*i.e*. replicates) were considered as random effects.

We used Generalized Linear Mixed-Effects Models (GLMMs) with Poisson distributions (*glmer* function, *lme4* package) to test how the number of flying mosquitoes (*n*_flying_) and swarming mosquitoes (*n*_swarming_) differs between swarming phases (start, peak and end phases). Finally, we used a binomial GLMM to test how the proportion of flying mosquitoes that swarm (*R*_swarming_ *= n*_swarming_*/n*_flying_×100%) differs between the swarming phases (start, peak and end phases). The experimental replicates were considered as random effects.

For model selection, we used the stepwise removal of terms, followed by likelihood ratio tests. Term removals that significantly reduced explanatory power (*P*<0.05) were retained in the minimal adequate model. Analyses for normality and homogeneity of variances were performed with a Shapiro and Fligner test, respectively (*shapiro.test* and *fligner.test* functions, *stats* package). If necessary, data were transformed using the Box-Cox power transformation method (*powerTransform* function, *car* package).

In addition, we performed analyses on the effect size. The value of partial Eta-squared (*η*^2^) was computed from the full models for each fixed effect (*eta_squared* function, *effectsize* package) and interpreted following the Cohen’s rule [50]. Thus, the effect of fixed terms was considered significant when it was both statistically significant (*P*<0.05) and with a medium or large effect size (*η*^2^>0.12). All statistical results are reported as mean±standard error or percentage [95% confidence interval].

## Results

### Flight activity during the swarming period

Mosquito flight behaviour exhibited a consistent pattern. As the ceiling lights dimmed, mosquitoes predominantly rested on the walls of the flight arena before initiating random flights approximately two minutes prior to lights-off (Figure 1*c*). Two to three minutes after the ceiling lights turned off, a male started to display a stationary looping flight over the marker. Then within 7-90 seconds (mean±se=46±8 s, *n*=5 experiments) additional mosquitoes joined the swarm, with the exception of a single replicate in which the first male swarmed alone for 233 seconds before departing the swarm location. The number of swarming mosquitoes steadily increased, reaching a swarming peak about 10 minutes later, during which the majority of flying individuals were engaged in swarming activity. This peak was relatively stable for approximately 20 minutes, after which the number of swarming mosquitoes gradually decreased back to zero (Figure 1*c*). It is worth noting that mosquitoes flying outside the swarm at the ending phase were flying mainly along the walls of the flight arena, bouncing against the walls as an apparent escape behaviour.

During the recordings, 18% to 73% of released mosquitoes showed flight activity, and 4% to 54% of them exhibited swarming behaviour. To systematically test how flight activity varied throughout the swarming period, we compared both the flight and swarming activity across the three swarming phases (start, peak and ending phase; Figure 1*c* and Table S1). This shows that both the number of flying mosquitoes and the number of swarming mosquitoes were statistically different between the three phases (Table S1). While the number of flying mosquitoes at the start and the end phases did not differ significantly, the number of flying mosquitoes was significantly higher during the peak phase compared to both the start and end phases (start: *n*_flying_=15.5±2.1 mosquitoes; peak: *n*_flying_=23.3±2.3; end: *n*_flying_=14.0±2.4; Table S1). Likewise, the number of swarming mosquitoes was significantly higher in the peak phase than in the other phases (start: *n*_swarming_=4.0±0.6 mosquitoes; peak: *n*_swarming_=16.7±2.1; end: *n*_swarming_=6.3±1.4; Table S1). As a result, the relative proportion of swarming mosquitoes to the total number of flying mosquitoes significantly differed across all phases, with the highest proportion observed during the peak phase, and the lowest at the start of swarming (start: *R*_swarming_=26% [9%]; peak: *R*_swarming_=71% [7%]; end: *R*_swarming_=45% [11%]; Table S1).

### Looping flight kinematics of individual swarming mosquitoes

The swarming male mosquitoes displayed a stereotypic stationary looping flight behaviour above the black ground marker (Figure 2). We summarised the looping flight kinematics of each flight track using the mean±standard error of the main kinematics parameters (Table S2): trajectory location, flight speed, acceleration, and the distance to nearest neighbour in the swarm.

We visualised positional kinematics of the looping flights using the median and quartile distributions for each trajectory (Figure 2*d,e*), and histograms of the location distributions (Figure 2*f*). These show that most mosquitoes swarms centred above the marker (*x̅*=0.00±0.00 m, *y̅*=0.00±0.00 m), at an average height of *z̅*=0.67±0.01 m (Table S2 and Figure 2*a-f*). The variation around this mean location due to the looping flight behaviour was significantly smaller in the direction parallel to the sunset horizon than that perpendicular to it (*σ̅*(x)=0.06±0.00 m and *σ̅*(*y*)=0.10±0.00 m, respectively; Table S2 and Figure 2*e,f*). The variation in height during the looping flights was significantly smaller to the variation in both the *x* and *y*-direction (*σ̅*(z)=0.04±0.00 m; Table S2 and Figure 2*d-f*).

The swarming mosquitoes flew at speeds ranging from 0.24 to 1.28 m/s (*U̅*=0.50±0.00 m/s), with accelerations ranging from 0.29 to 7.33 m/s^2^ (*A̅*=2.88±0.02 m/s^2^). The average distance to the nearest neighbour was *d̅*_neighbour_=0.11±0.00 m.

The three flight speed components (*u,v,w*) were significantly different from each other, with the mean vertical speed (*w*) being the lowest and the mean horizontal speed normal to the sunset horizon (*v*) the highest (Table S2). The three flight velocity components showed distinct distributions (Figure 2*g*). Particularly, the flight velocity component normal to the sunset horizon (*v*) showed a distinctly wider double-peak distribution. As a result, the mean flight speed in this direction was 1.6 and 4.8 times higher than the flight speed parallel to the sunset horizon and the vertical speed, respectively (*u̅*=0.24±0.00 m/s, *v*=0.38±0.00 m/s and *w*=0.08±0.00 m/s; Table S2). Similar to the flight speed, the mean acceleration was highest in the direction normal to the sunset horizon, as it was on average 1.2 and 3.5 times higher than those parallel to the sunset horizon and vertically, respectively (*A̅*_*x*_=1.57±0.01 m/s^2^, *A̅*_*y*_=1.90±0.02 m/s^2^, *A̅*_*z*_=0.55±0.00 m/s^2^; Table S2).

We also analysed the interaction between the three-dimensional axes and swarming phases (axes × phases interactions; Table S4) to test whether the *x*, *y* and *z* components of flight kinematics parameters varied over the three swarming phases. For all the tested looping flight kinematics parameters we found no significant axes × phases interaction (Table S4), meaning that the three components (*x*,*y*,*z*) of the looping flight kinematics parameters were consistent over the three phases. Furthermore, we found that the tested looping flight kinematics parameters were consistent among the replicates (location: 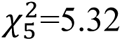, *P*=0.378; Figure S5).

We consequently tested whether the swarming kinematics differed between the three swarming phases (Table S3). Most of the kinematics parameters did not differ significantly among the phases, including the mean and standard deviation of the looping trajectory locations, the mean speed and mean acceleration during the looping flights (Table S3). Only the distance to the nearest neighbour significantly differed across phases: at the peak phase, the mean distance to the nearest neighbour was 1.9 and 1.2 times smaller than at the start and end phases, respectively (start: *d̅*_neighbour_=0.19±0.01 m; peak: *d̅*_neighbour_=0.10±0.00 m; end: *d̅*_neighbour_=0.12±0.00 m; Table S3).

Finally, we tested how the swarming kinematics varied with the swarm size at the swarming peak phase when the swarm size was relatively consistent (Table S3 and Figure S6). Again, only the distance to the nearest neighbour varied significantly with swarm size, as it decreased with increasing swarm size with a rate of 2 mm per additional swarming mosquito (Figure S6*e*).

### Size, shape and structure of the complete swarm

By combining the individual swarming flight tracks, we quantified the emerging kinematics of the swarm as a whole, for the three swarming phases separately (Figure S3), for all the six replicates separately (Figure S4), and for all swarming mosquitoes combined (Figures 3 and 4). For both the replicates and the three swarming phases, we visualised the swarming kinematics using the spatial distribution of normalised swarming density (Figures S3 and S4). We observed that the size, shape and structure of swarms were consistent over the phases and across the six replicates (Figures S3 and S4, respectively). This similarity across phases and replicates allowed us to combine all swarming data into a large dataset in order to reconstruct the average swarm, with which we subsequently worked (Figures 3 and 4).

The observed swarm is shaped as a flattened elliptical cone, because it is wider at the top (∼0.5 m) than at the bottom (∼0.2m) (Figures 2*a-e* and 3*a-c*), and it is slightly stretched horizontally in the direction of the sunset horizon (*y*-axis). The centre of the swarm is positioned straight above the visual ground marker, and the top and bottom of the swarm were located at a height of approximately 0.95 m and 0.35 m, respectively (Figures 2*a* and 3*a-c*).

The density of swarming flight points and tracks was higher in the middle of the swarms (Figure 3*d-i*) gradually decreasing towards top, bottom and side boundaries. Particularly, at the top and bottom regions, densities were approximately three times lower than in the middle section (Figure 3*d,g*). Nevertheless, mosquitoes that went in the lower half of the swarm stayed significantly longer in this region with durations approximately twice of those in the upper section (Figure 3*j-l*). In contrast to the density of swarming flight tracks, the distance to the nearest neighbours was the lowest in the middle of the swarm with mosquitoes flying around two times as close to each other in the middle of the swarm than at boundaries (Figure 4*a-c*).

Examining the spatial distributions of the flight kinematics parameters reveals that swarming mosquitoes were flying 1.25 times faster in the top half of the swarm (Figure 4*d-f*) compared to the lower section (Figure 4*d-f*). However, the accelerations were the highest at the boundaries of the swarm, gradually reducing to nearly zero along the central axis of the swarm *y*=0 m (Figure 4*g-i*).

### Modelling mosquito swarming behaviour

After finding that the swarm kinematics were highly consistent across the three swarming phases and replicates, we used the average swarm location, size and shape to develop, test and validate our sensory-cue informed swarming model (Figures 5 and S12). We first developed a mathematical representation of the average swarm reconstructed from our experimental data (Figure 5*b,d-f*). This consists of the previously described flattened elliptical cone, of which the short and long primary ellipse axes increase in size with height (Figures 5*b* and S12*a,d*). The long axis is oriented normal to the sunset horizon, and the short axis is oriented parallel to it.

Our swarm model assumes that swarming mosquitoes remained within this swarm volume by using visual cues, particularly optical angle of the ground marker (Figures 5*a* and S12*b,c*). We therefore calculated the optical angles that mosquitoes would perceive of the marker throughout the complete swarming area (Figure 5*d-f*). Because the swarm circumference is elliptical, we calculated the optical angle of the marker along the *x* and *y*-axis separately (*α* and *β*, respectively; Figure S12*b,c*). Optical angles of an object are inversely proportional to the distance from that object, and thus the optical marker angles decrease with increasing height above the marker, but also when moving horizontally from the middle of the marker to the edges.

We then used these optical data in combination with the mathematical representation of the average swarm to estimate how swarming mosquitoes could use their visual perception of the ground marker to remain within the swarm volume. We defined the top and bottom boundaries of the swarm based on absolute optical angle values (Figure 5*d,e*), and the cone-like side edges of the swarm based on the change in optical angle (optic flow cues).

We first tested how well the optical angles of the marker in the directions parallel and normal to the sunset horizon captured the observed elliptical swarm shape (Figure S12*b,c*). For this, we calculated both optical marker angle parameters *α* and *β*, at the swarm edge along both the short and long ellipse axes (*x* and *y* directions). This showed that, for any given height, the *α* optical angles computed at the edge of the swarm are the same along both axes (Figure S12*b*), but they differ for the *β* optical angles (Figure S12*c*). This suggests that swarming mosquitoes use the optical angle parallel to the sunset horizon (*α* in *x*-direction) to remain within the elliptical circumference of the swarm.

Secondly, we tested what might cause the increase in swarm circumference with increasing height. We hypothesized that swarming mosquitoes use the change in optical angle as cue to stay inside the swarm edge and move back towards the middle. To test this, we determined at various heights the percentage decrease in the *α* and *β* optical marker angles when moving from the swarm centre towards the edge, along both ellipse axes (Figure S12*e,f*). The relative optical angle change along the *x*-axis is defined as *R*_α_=(α_max_-α)/α_max_×100%, where α_max_ is the maximum optical angle at the centre of the swarm in the *x*-direction. The relative optical angle *R*_α_ at the edge in the *x*-direction turned out to be consistently 4.5% of the maximum angle, along both ellipse axes and at all swarm heights (Figure S12*e*). In fact, the 4.5% threshold at a given height exactly produces the elliptical circumference observed in the swarming experiments (Figure 5*f*,*i*). In contrast, the relative optical angle *Rβ* at the edge in the *y*-direction differed between both ellipse axes, as it was consistently 2.5 and 8.0 along the *x* and *y*-axis, respectively, at all swarm heights (Figure S12*f*). This further supports the notion that swarming mosquitoes use the optical angle in the *x*-direction (*α*) to remain within the circumference of the swarm.

Because swarming mosquitoes apparently use the optical angle of the marker in the *x*-direction (*α*) to control their horizontal position within the swarm, we determined the optical angle *α* at the bottom and top of the swarm (Figure 5*d*,*e*). It appears that at the bottom and top of the swarm mosquitoes experienced an *α* optical angle of the marker of 55° and 24°, respectively. These angles could thus be used by mosquitoes as thresholds to conform to the swarm height distribution. The resulting modelled swarm shape and size captured the experimentally measured swarm well (Figure 5*b-i*).

## Discussion

### Mosquito flight activity during swarming

By combining videography-based experiments with modelling, we studied the flight kinematics of *An*. *coluzzii* male mosquitoes during simulated sunset and how their collective flights result in emergent swarming behaviour. At the start of sunset, resting males first started to fly throughout the complete flight arena. Then, the swarm was initiated by a male performing a stationary pseudo-looping flight over a visual ground marker. Within the following minutes, it was sequentially joined by other mosquitoes, resulting in the development of a swarm. The number of swarming males increased over time to reach a peak swarming phase approximately ten minutes after the start. The swarm remained relatively stable for approximately ten minutes, after which the swarm size gradually decreased, and swarming ended approximately thirty minutes after it started. The mosquitoes that stopped swarming often kept flying. This post-swarming flight activity is distinctly different from the pre-swarming activity, as here the male mosquitoes repeatedly fly against the inner walls of the arena.

On the assumption that both male and female mosquitoes use swarm markers to find and join mating sites [20,21,23], we suggest that at the start of sunset, the non-swarming flight continues until the mosquito find an appropriate visual ground marker. Then, the male mosquito initiates a stationary pseudo-looping flight above the marker until it detects a potential mate. The presence of swarming male mosquitoes at a site may also visually attract other males to the swarm, but this remains to be proven. The post-swarming flight activity near the walls of the arena suggests that these mosquitoes exhibit a dispersal flight behaviour, possibly seeking for a food source after their energetically costly swarming flights [51,52].

During our experiments, we simulated sunset light conditions by slowly reducing ceiling light intensity and providing a sunset horizon. The increase in flight activity and the start of swarming were both triggered by the sunset simulation, although the circadian clock also plays a key role in the process [41,53,54]. After the initial sunset simulation, light intensity in the experimental room were kept constant. Thus, light conditions remained unchanged from 2-3 minutes before the first male started swarming until after the ending of swarming. Because swarm ending occurred approximately 30 minutes after swarm initiation, swarm ending could not have been triggered by changes in light conditions. Swarming flight behaviour has an exceptionally high energetic cost [51,52], suggesting that the male mosquitoes stopped swarming when a threshold in their energy reserve was reached.

Therefore, the post-swarming flight activity near the flight arena walls could be the result of an escape-like flight behaviour or straight flight of mosquitoes in search for a sugar meal to refuel their reserves. If energy reserves limit the duration that a male mosquito is able to remain swarming, then presence in a swarm is a good signal of male fitness for the female seeking a mate [55]. Swarming duration could thus equally be used by researchers to assess the potential fitness of laboratory male mosquitoes used in mosquito release programs [21,23,41,56].

### Location and shape of the swarm and viewing angle of the swarm marker

As expected in *An. coluzzii* mosquitoes, males located their swarms above the visual ground marker. This behaviour has been previously reported and discussed as the primary premating barrier governing the swarm spatial segregation and the reproductive isolation in *An. Gambiae s.l*. [18,21,57].

The swarm was consistently located right above the marker at an average height of 0.67 m and was shaped like an upside-down elliptical flattened cone. The swarming mosquitoes flew above the marker with optical viewing angles of the marker ranging from 24° to 55°. These values would be the minimum and maximum thresholds in our experimental conditions to set the height range of the swarm above the marker. If true, a larger marker should induce higher and wider swarms, which was reported by Poda et al. [21]. This would suggest that these features are probably governed by environmental visual cues, but the consistency of viewing angle range with different marker sizes needs to be confirmed in further studies.

As both sexes use the marker to join the species-specific mating site [17,21,24], the range of viewing angle used by both males and females need to be consistent and species-specific to bring sexes together. This relation of the swarm to the marker was also closely studied in other mosquito species. In laboratory conditions, *Culex pipiens quinquefasciatus* males formed swarms above a 26 cm square marker where the marker viewing angle ranged from 40° to 90° [24]. However, the height of swarms may differ locally [58] and consequently, the marker viewing angle could depend on local adaptations.

The upside-down cone shape of the swarm can be explained by the fact that at a given height, males did not deviate in the horizontal plane by more than 4.5% from the maximum viewing angle. As a result, the horizontal flight amplitude that allows staying within the swarm volume increases with height. Our mathematical model is based on the optical angle of the marker and predicted well the location and shape of the swarm found from the experimental data, assuming that these are primarily based on optical metrics of the swarm marker.

A similar swarm shape has been observed by Cavagna et al. [32] in laboratory male swarms of *An. coluzzii*, while various other swarm shapes have been observed in the field [26,59]. These differences in swarm shape could be due to variations in the shapes of the visual markers used [26,59]. In addition to the marker features, the surrounding skyline and the wind speed and direction could also influence the swarm characteristics [17,26,60].

### Structure of the swarm and spatial dynamic of flight kinematics in the swarm volume

Within the swarm volume, mosquitoes flew mainly in the horizontal plane with variable flight speeds and accelerations. Both horizontal flight speeds and accelerations were higher in the direction of the artificial sunset horizon (*y*-axis) than parallel to it (*x*-axis). This asymmetric pattern in the flight kinematics resulted in the flattened elliptical cone shape of the swarm with the major ellipse axis in the direction of sunset horizon (*y*-axis). This could be due to the location of the marker in the flight arena and/or to the rectangular shape of the flight arena. However, the widest side of the flight arena was parallel to the sunset horizon (*x*-axis), which is the direction in which the swarm was narrowest. Furthermore, this flight pattern has also been reported in natural swarms of *An. gambiae s.l.* (*i.e.* east–west flight direction) [30].

The flattened elliptical swarm shape is thus more likely to be a behavioural orientation in relation to the sunset rather than a set-up bias. This is further supported by our swarming model that suggests that our swarming mosquitoes consistently use the optical marker angle parallel to the sunset horizon for positioning themselves above the marker. Orienting towards the sunset would not only allow swarming mosquitoes to robustly assess this optical angle of the marker, but also generate a good viewing of the surrounding and consequently of potential mates entering the swarm.

In contrast to the primarily horizontal flight pattern of swarming *An. gambiae s.l.*, *Cx. pipiens quinquefasciatus* and *Aedes albopictus* exhibit mainly vertical flight movements in swarms (Gibson [24] and Gillies (personal communication), respectively). This suggests that our results could be specific to *An. gambiae s.l.* [21,30,32].

Our recordings revealed a higher density of flight tracks in the middle of the swarm resulting in shorter distances between individuals in this region, something that was also reported from natural swarms of *An. gambiae s.l*. [26,30,31]. According to Manoukis et al. [26], this location could confer an advantage to the males as it allows a quicker access to any part of the swarm periphery and thus a quicker access to females if they randomly enter swarms. In addition, once in the swarm, females may simply be more likely to pass through the middle of the swarm, making this location a hotspot for mating. Alternatively, this location in the swarm could simply be a by-product of mosquito perception of the marker. Our results provide further support for the latter hypothesis. Indeed, if males must remain directly above the marker without losing eye-contact with it and have to fly within a given range of marker viewing angle, which probably contains an optimal value, a pattern such as the one observed could emerge. The middle of the swarm could be the region of the optimal marker viewing angle. If this hypothesis is true, the same swarming flight pattern will be observed in any species using a fixed visual landmark to define the location and volume of the swarm.

Another hotspot could be the lower half of the swarm, as we found that males spent more time in this area. Butail et al. [30] reported that females were observed predominantly in this area just before the copula. This location in the swarm could also simply be a by-product of the perception of the marker by both the male and female mosquitoes.

The spatial distributions of flight kinematics parameters show that swarming mosquitoes were flying faster in the top half of the swarm, and the flight accelerations were higher at the boundaries and close to zero along the central axis of the swarm. This is probably due to the swarm shape and its fixed volume, both governed by the mosquito perception of the marker. First, swarming individuals were mainly flying horizontally and remained in the area above the marker by flying at relatively constant speeds in the centre of the swarm and rapidly turning at the swarm edges, leading to high accelerations. Secondly, the upside-down cone shape of the swarm provides large horizontal flight amplitudes in the top region of the swarm, allowing faster flights in this region. This could explain the fact that mosquitoes spent less time in this region because of the high energetic cost of the swarming flight [51,52].

### Temporal dynamic of the swarm structure and flight kinematics of individual swarming mosquitoes

Except for the number of individuals in the swarm and the distance to the closest neighbour, the characteristics of the swarm as a whole did not significantly change between the start, peak and ending phases of swarming activity. The location, dimensions, and shape of the swarm as well as the flight pattern of the swarming individuals were consistent over the three swarming periods, regardless of the number of individuals in the swarm. This has also been reported in *Culex pipiens pallens* [25], and supports the hypothesis that the location, volume, and shape of the swarm are defined by the visual characteristics of the marker [21,24,26,59]. However, Feugère et al. [23] showed differences between *An. coluzzii* and *An. gambiae s.s.* in radius of field swarms as a function of number of males, where *An. gambiae s.s.* showed a positive effect of number of swarming males over the swarm radius. Whatever the case, the swarming space, which would be defined by the marker size, cannot contain an infinite number of individuals due to a saturation effect in the swarm size [21]. Each male in a swarm would need a given airspace to produce the stereotyped swarming flight.

While the volume of the swarm remained consistent, the individuals in the swarm varied over the swarming period, resulting in a variation in the distance to the nearest neighbour over the swarming period. It decreased during the peak phase corresponding to the highest swarm density observed during this period, and was negatively correlated to the number of individuals in the swarm at the peak of swarming. However, swarming individuals need to keep some minimal distance from each other to display the stereotypic swarming flight while avoiding collusions; hence, the saturation effect in swarm size reported by Poda et al. [21].

Regarding the mosquito flight kinematics, neither the flight speed nor acceleration changed significantly between the three swarming phases. This suggest that male mosquitoes in the swarms have the same ability to mate over the swarming period, and mating can successfully occur within the swarm at any time of the swarming period.

### Mosquito swarming in a broader context

Our experimental set-up allowed three-dimensional recording of the flight activity of *An. coluzzii* males at the start, peak and ending phase of swarming. Data analysis provided relevant quantitative information on the spatial and temporal dynamics of the swarm characteristics, mosquito flight kinematics and their relationship with the swarm marker. All these metrics were highly consistent across replicates and the three swarming phases suggesting the repeatability of the experiments, and the consistency of the swarming individual behaviour over the swarming period, respectively. However, the swarm had a non-homogeneous structure and a differential spatial distribution of the swarming flight kinematics, most likely governed by the mosquito perception of the swarm marker.

These findings may help to explain mechanisms underlying speciation and diversification within the *An. gambiae* complex as any single significant change in the swarming flight behaviour or in the utilization of the swarm marker could lead to divergence in a mosquito population and promote speciation. Similar works within the *An. gambiae* complex may provide information on which of the studied metrics differ between the species and thus can be involved in speciation or conspecific recognition.

In the field of vector control, such changes may be induced during colonisation or genetic modification of mosquitoes used in release programmes potentially compromising their competitiveness and mating success in the field. Therefore, these metrics may be used to assess impact of colonisation or genetic modifications on mosquito swarming behaviour, or to gauge competitiveness of produced males.

## Acknowledgements

We thank Somda Stephane, Sougué Emmanuel, Guinko Nourou, Coulibaly Valérie and Bandaogo Abdoul Malik for mosquito rearing. We are grateful to Wageningen University and Research for training and material support during the data analysis and drafting of the manuscript.

## Authors’ contributions

B.S.P., O.R., M.F., P.M. installed and calibrated the video tracking system. B.S.P., O.R. conceived the experimental design. B.S.P., C.N., D.F.S.H. performed data collection. B.S.P., A.C., F.T.M. performed data analyses. B.S.P., A.C. drafted the figures. B.S.P., M.F., L.F. performed data analyses on a preliminary experiment. B.S.P. drafted the manuscript and A.C., F.T.M., O.R., L.F. critically revised the manuscript. All authors revised the manuscript, gave final approval for publication and are accountable for the work performed therein.

## Ethics approval and consent to participate

Not applicable.

## Supplementary information

The online version contains supplementary material available from…

## Availability of data and materials

All original Matlab and R codes as well as the analysed three-dimensional tracking data of swarming mosquitoes is publicly available in the DRYAD repository: https://datadryad.org/stash/share/oj0wQHGLL68s3h8mm7fJog3XdqFyuJCpetK9LqGuyUw. The raw datasets are available from the corresponding author upon reasonable request.

## Declaration of AI use

We have not used AI-assisted technologies in creating this article.

