## Supplementary File for "Spatial and temporal characteristics of laboratory-induced *Anopheles coluzzii* swarms: shape, structure and flight kinematics"

### Supplemental figures

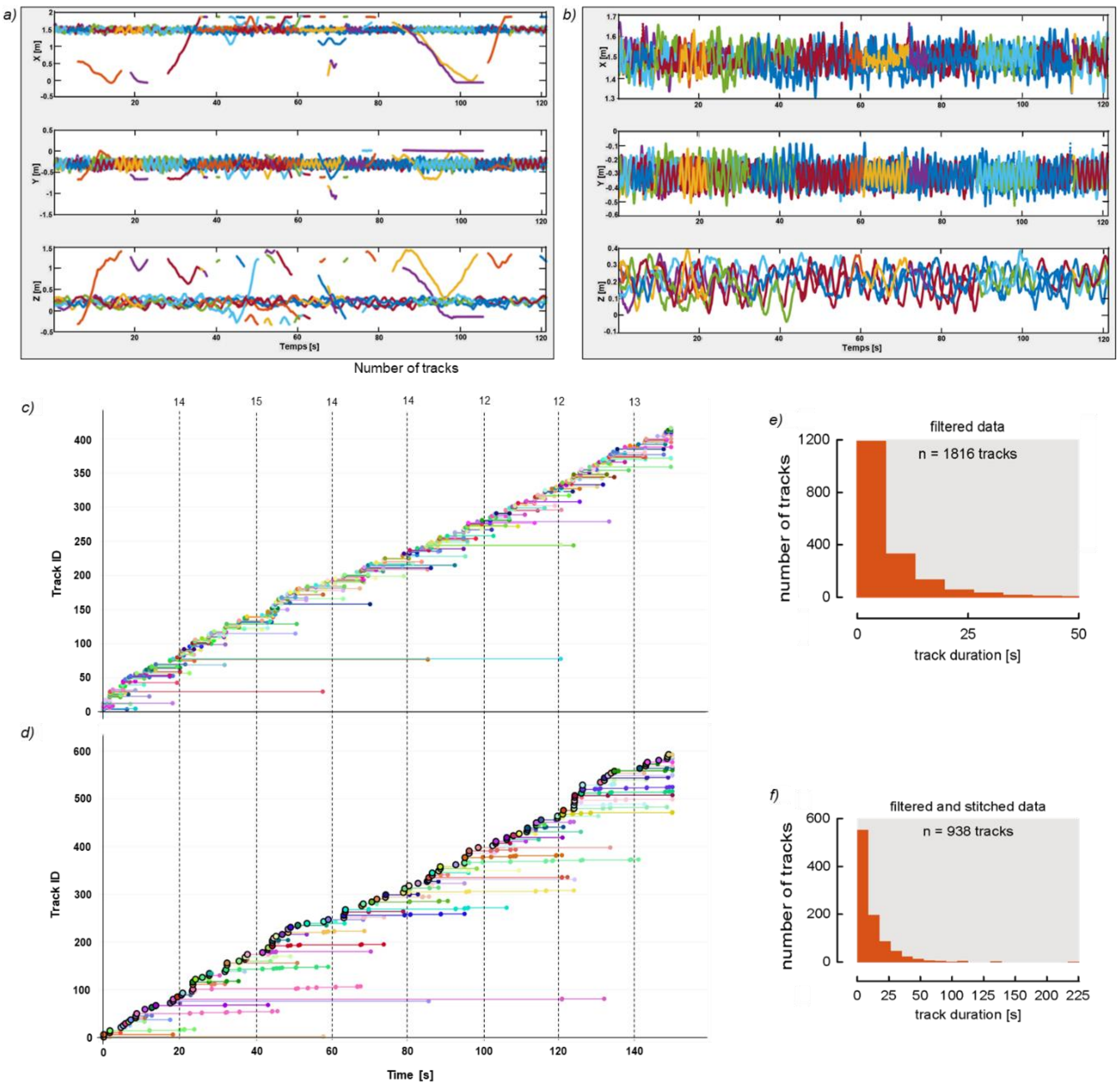


**Figure S1: Selection of swarming tracks and reconstruction of interrupted trajectories of swarming mosquitoes.** (*a*) Flying tracks plotted as the instantaneous three-dimensional locations of mosquitoes and (*b*) sinusoidal swarming tracks manually selected using a MATLAB program (Data Selector, SciTrackS Gmbh, Bertschikon, Switzerland). (*c*) Selected swarming tracks (*b*) automatically combined using a R program (RMosquito, SPILEn, France). These graphs (*a*-*d*) show the data of the swarm #2 recorded at the peak period of the swarming activity. (*e*,*f*) Histogram of the track duration of (*e*) the selected swarming tracks showing that the duration of the raw tracks mainly ranged between 0 and 5 seconds and (*f*) combined swarming trajectories of all the recordings showing that most swarming mosquitoes remained in the swarm for at least 25 seconds. Indeed, for the six experimental replicates, we reconstructed in total 13,474 flight tracks. These tracks were from both flying non-swarming mosquitoes and mosquitoes that exhibited distinct swarming behaviours above the ground marker. (*e*) Manual selection resulted in the identification of 1,816 swarming tracks, and (*f*) consecutive automatic stitching of these swarming tracks resulted in a final dataset of 938 swarming flight trajectories. The duration of these swarming flight tracks varied from a couple of seconds for the short tracks, to a maximum of 233 seconds for the longest track. Of these 938 swarming flight trajectories, 60 occurred during the swarm start phase, 705 during the swarm peak phase, and 173 during swarm ending.

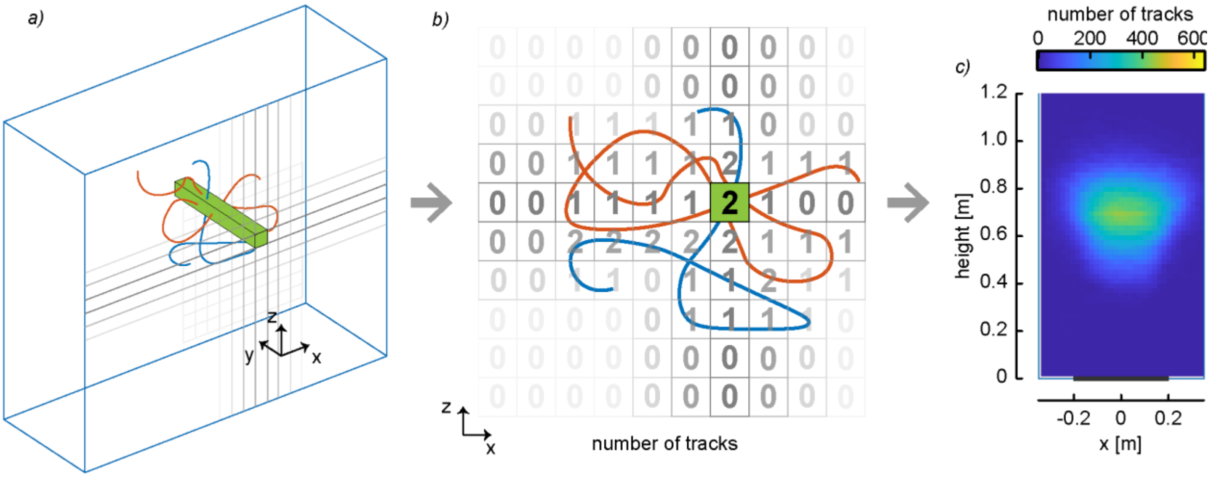


**Figure S2: Analysis method to visualize mosquito spatial flight dynamics.** (*a*) The recorded volume is divided in smaller rectangular sub-volumes of 2 cm × 2 cm × depth. (*b*) Metrics (*e.g.* number of tracks) are computed for each sub-volume. (*c*) These metrics are then visualised in two-dimensional heatmaps. Here we can see the spatial distribution of the number of mosquito tracks for all swarms.

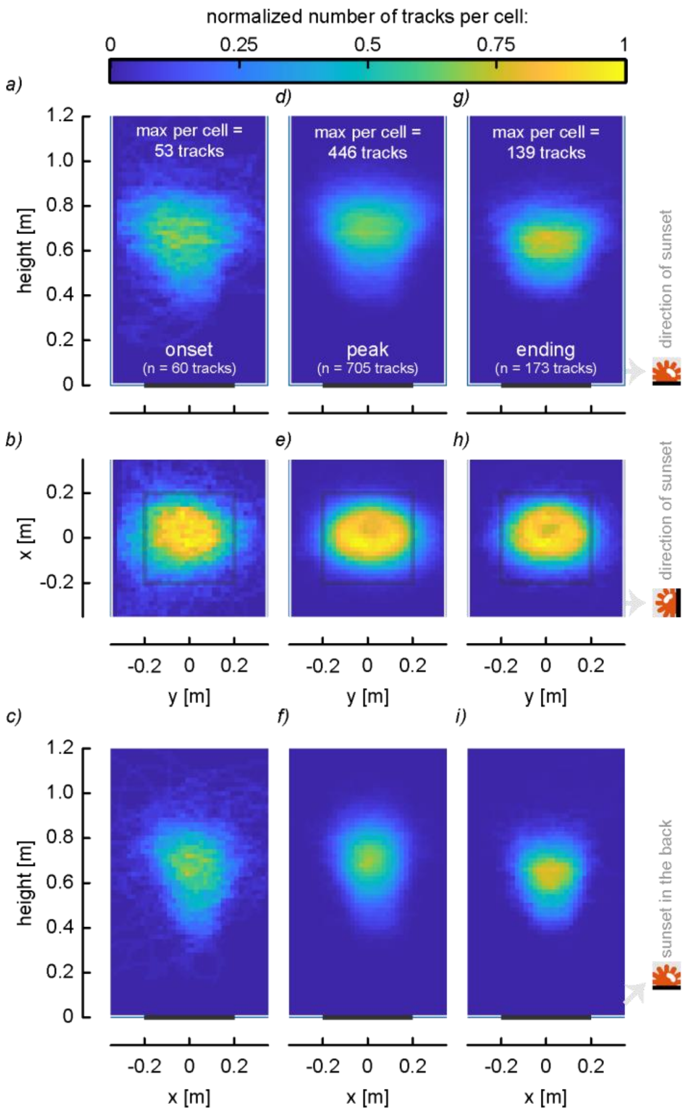


**Figure S3: Swarm shape and spatial distribution of swarming track density at the three swarming phases.** The spatial distribution of the number of swarming tracks per cell recorded at (*a*-*c*) the start, (*d*-*f*) peak and (*g*-*i*) ending phases of swarming activity. The number of tracks for each swarm has been normalized by the maximum number of tracks computed per cell in order to facilitate comparison between swarming phases. Depending on the viewing plane, the swarm marker is represented by a black square or a black line.

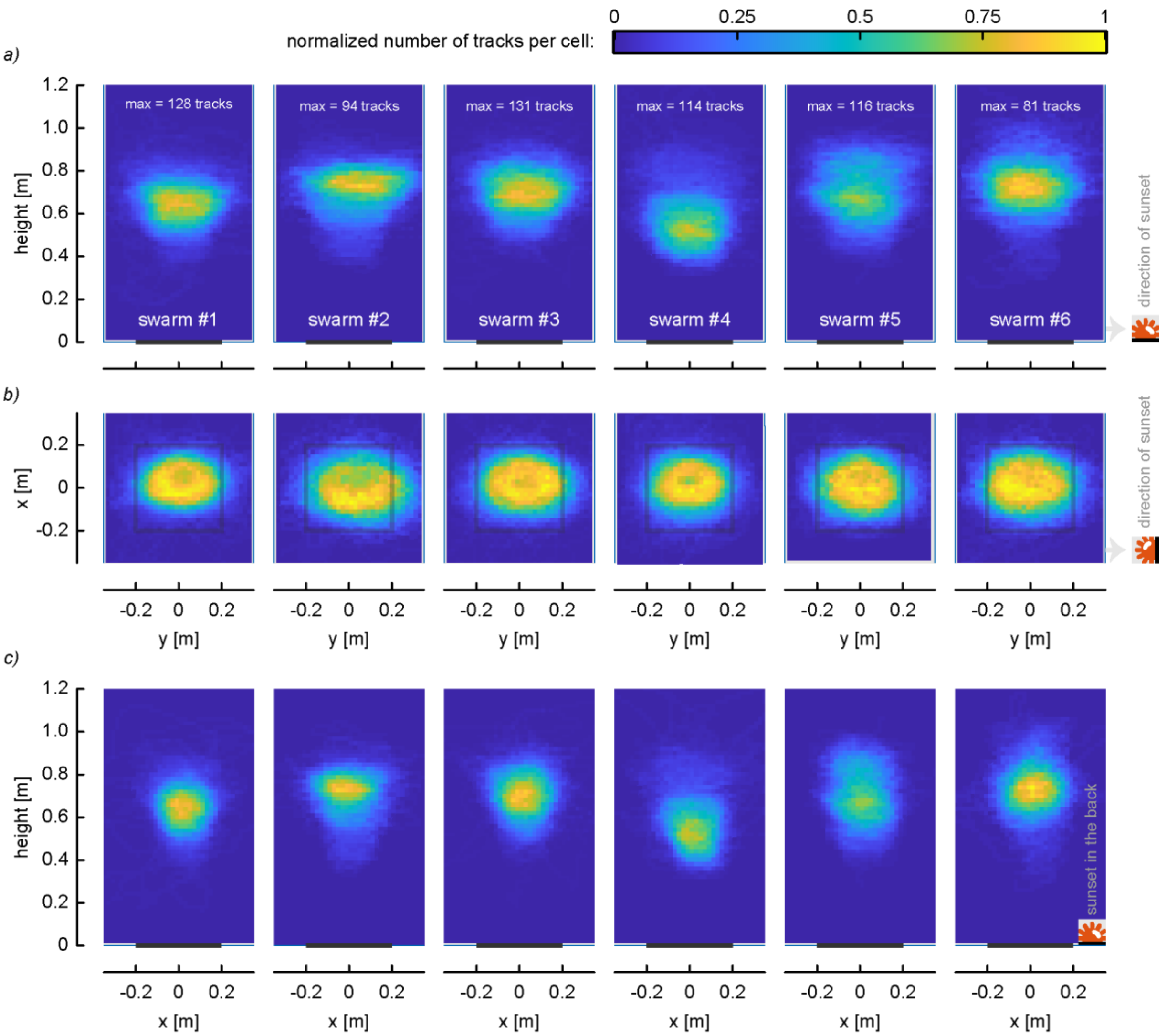


**Figure S4: Spatial distribution of track density for each replicate.** (*a*) Side view, (*b*) top view and (*c*) front view of the spatial distribution of the number of swarming tracks for each of the six swarms. Depending on the viewing plan, the swarm marker is represented by (*b*) a black square or (*a*,*c*) a black line. The number of tracks for each swarm has been normalized by the maximum number of tracks computed per 2 cm × 2 cm cell in order to facilitate comparison between swarming events. We can observe a reproducibility of the swarm shape and structure during the experiments.

**Table S1:** Results of the Generalized Linear Mixed Models testing how the flight and swarming activity varied between the three swarming phases (start, peak and end of swarming).

| | | **Swarming phases** | | | | | | |
|---|---|---|---|---|---|---|---|---|
| | | **Start** | **Peak** | **End** | **$X^2$** | **df** | **P-value** | **$\eta^2$ [Interpretation]** |
| **Number of flying mosquitoes** | **[Min - Max]** | [9 - 22] | [19 - 34] | [8 - 24] | | | | |
| $n_{flying}$ | **Mean ± se** | 15.5 ± 2.1 *a* | 23.3 ± 2.3 *b* | 14.0 ± 2.4 *a* | 16.80 | 2 | **< 0.001** | **0.39 [large]** |
| **Number of swarming mosquitoes** | **[Min - Max]** | [2 - 6] | [13 - 27] | [3 - 11] | | | | |
| $n_{swarming}$ | **Mean ± se** | 4.0 ± 0.6 *a* | 16.7 ± 2.1 *b* | 6.3 ± 1.4 *a* | 53.32 | 2 | **< 0.001** | **0.73 [large]** |
| **Swarming percentage** | **[Min - Max]** | [20% - 33%] | [65% - 79%] | [23% - 62%] | | | | |
| $R_{swarming} = n_{swarming}/n_{flying} \times 100\%$ | **Percentage [95% CI]** | 26% [9%] *a* | 71% [7%] *b* | 45% [11%] *c* | 44.43 | 2 | **< 0.001** | - |

se: standard error. CI: confidence interval. Different letters indicate statistically significant differences between swarming phases at $P < 0.001$. Cases with both statistically significant differences ($P < 0.001$) and large effect size ($\eta^2 > 0.12$) are in bold.

**Table S2**: Results of the Mixed Linear Models testing whether there were variations between the *x*, *y* and *z* components of the various flight kinematics parameters.

| | | Three-dimensional axes | | | | | | |
|---|---|---|---|---|---|---|---|---|
| | | ***x*-axis** (parallel to sunset horizon) | ***y*-axis** (normal to sunset horizon) | ***z*-axis** (height) | **$X^2$** | **df** | **P-value** | **Partial $\eta^2$ [Interpretation]** |
| **Location (m)** | **[Min - Max]** | [-0.16 - 0.22] | [-0.27 - 0.30] | [0.30 - 0.93] | | | | |
| | **Mean ± se** | 0.00 ± 0.00 *a* | 0.00 ± 0.00 *a* | 0.67 ± 0.00 *b* | 1100.68 | 2 | **< 0.001** | **0.89 [large]** |
| **Standard deviation to the mean location (m)** | **[Min - Max]** | [0.01 - 0.15] | [0.01 - 0.21] | [0.00 - 0.14] | | | | |
| | **Mean ± se** | 0.06 ± 0.00 *a* | 0.10 ± 0.00 *b* | 0.04 ± 0.00 *c* | 81.06 | 2 | **< 0.001** | **0.24 [medium]** |
| **Velocity (m/s)** | **[Min - Max]** | [0.04 - 0.50] | [0.07 - 1.22] | [0.02 - 0.21] | | | | |
| | **Mean ± se** | 0.24 ± 0.00 *a* | 0.38 ± 0.00 *b* | 0.08 ± 0.00 *c* | 846.85 | 2 | **< 0.001** | **0.69 [large]** |
| **Acceleration (m/s²)** | **[Min - Max]** | [0.07 - 4.13] | [0.15 - 6.01] | [0.07 - 2.35] | | | | |
| | **Mean ± se** | 1.57 ± 0.01 *a* | 1.90 ± 0.02 *b* | 0.55 ± 0.01 *c* | 343.48 | 2 | **< 0.001** | **0.48 [large]** |

se: standard error. Different letters indicate statistically significant differences between swarming periods at $P < 0.001$. Cases with both statistically significant differences ($P < 0.001$) and medium or large effect size ($\eta^2 > 0.12$) are in bold.

**Table S3**: Results of the Mixed Linear Models used to test how the various flight kinematics parameters varied between the three swarming phases (start, peak and end of swarming).

| | | Swarming phases | | | | | | | Swarm size | | | |
|---|---|---|---|---|---|---|---|---|---|---|---|---|
| | | **Start** | **Peak** | **End** | **$X^2$** | **df** | **P-value** | **Partial $\eta^2$ [Interpretation]** | **$X^2$** | **df** | **P-value** | **Partial $\eta^2$ [Interpretation]** |
| **Height (m)*** | **[Min - Max]** | [0.30 - 0.84] | [0.43 - 0.93] | [0.44 - 0.92] | | | | | | | | |
| | **Mean ± se** | 0.62 ± 0.01 *a* | 0.68 ± 0.00 *b* | 0.62 ± 0.01 *a* | 48.61 | 2 | < 0.001 | 0.05 [small] | 0.70 | 1 | 0.403 | 0.001 [very small] |
| **Standard deviation to the mean location (m)** | **[Min - Max]** | [0.07 - 0.24] | [0.03 - 0.22] | [0.03 - 0.19] | | | | | | | | |
| | **Mean ± se** | 0.15 ± 0.00 *a* | 0.13 ± 0.00 *b* | 0.12 ± 0.00 *c* | 34.69 | 2 | < 0.001 | 0.04 [small] | 0.73 | 1 | 0.392 | 0.004 [very small] |
| **Velocity (m/s)** | **[Min - Max]** | [0.25 - 0.69] | [0.24 - 1.28] | [0.32 - 0.80] | | | | | | | | |
| | **Mean ± se** | 0.45 ± 0.01 *a* | 0.52 ± 0.00 *b* | 0.48 ± 0.00 *c* | 68.22 | 2 | < 0.001 | 0.07 [small] | 0.212 | 1 | 0.645 | 0.0004 [very small] |
| **Acceleration ($m/s^2$)** | **[Min - Max]** | [1.13 - 4.65] | [0.29 - 7.33] | [0.95 - 5.67] | | | | | | | | |
| | **Mean ± se** | 2.88 ± 0.10 *a,b* | 2.92 ± 0.03 *a* | 2.72 ± 0.05 *b* | 10.53 | 2 | < 0.01 | 0.01 [very small] | 0.322 | 1 | 0.570 | 0.001 [very small] |
| **Distance to nearest neighbour (m)** | **[Min - Max]** | [0.08 -0.35] | [0.03 - 0.23] | [0.06 - 0.30] | | | | | | | | |
| | **Mean ± se** | 0.19 ± 0.01 *a* | 0.10 ± 0.00 *b* | 0.12 ± 0.00 *c* | 337.4 | 2 | **< 0.001** | **0.27 [large]** | 133.34 | 1 | **< 0.001** | **0.22 [medium]** |

*: As the swarm was always located above the marker, the swarm location analysed here was the height. se: standard error. Different letters indicate statistically significant differences between swarming periods at $P < 0.01$. Cases with both statistically significant differences ($P < 0.01$) and medium or large effect size ($\eta^2 > 0.12$) are in bold.

**Table S4**: Effect of the 'axes × swarming phases' interaction on the swarming mosquito location and flight kinematics.

| | Axes × Swarming phases | | | |
|---|---|---|---|---|
| | $X^2$ | df | P-value | Partial $\eta^2$ [Interpretation] |
| **Location (m)** | 11.41 | 4 | 0.022 | 0.01 [very small] |
| **Standard deviation to the mean location (m)** | 46.90 | 4 | < 0.001 | 0.02 [small] |
| **Velocity (m/s)** | 46.44 | 4 | < 0.001 | 0.01 [very small] |
| **Acceleration (m/s²)** | 19.62 | 4 | < 0.001 | 0.006 [very small] |

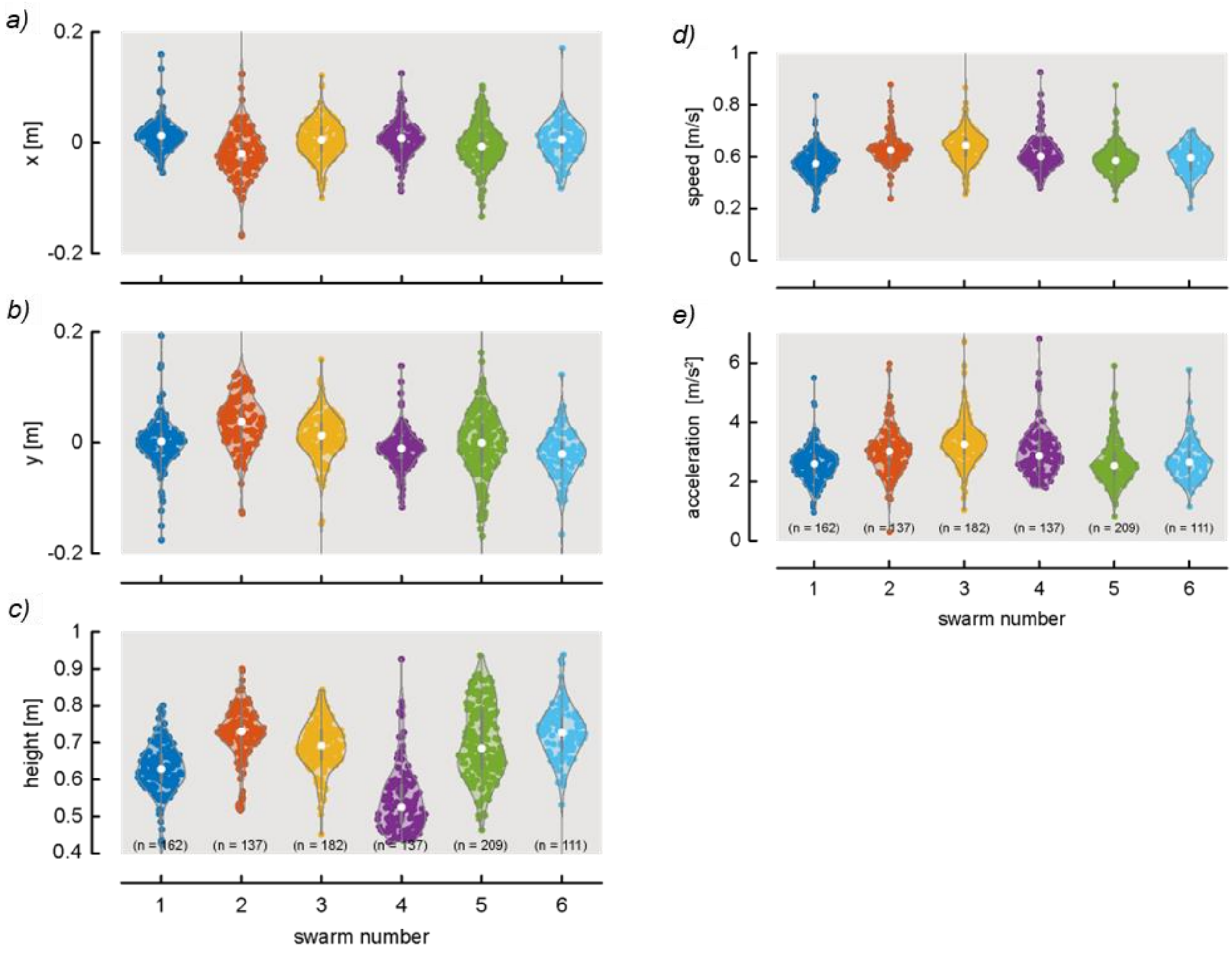


**Figure S5: Three-dimensional locations and flight kinematics of swarming mosquitoes in each recorded swarm**. (*a*-*c*) Violin plots showing the mean location of swarming mosquitoes along the *x*, *y* and *z*-axis. (*d*,*e*) The mean flight speed and acceleration of swarming mosquitoes in each swarm. *n* equals the number of tracks for the specific swarming experiment (replicate). We can observe a reproducibility of the swarm locations and flight kinematics of swarming mosquitoes during the experiments.

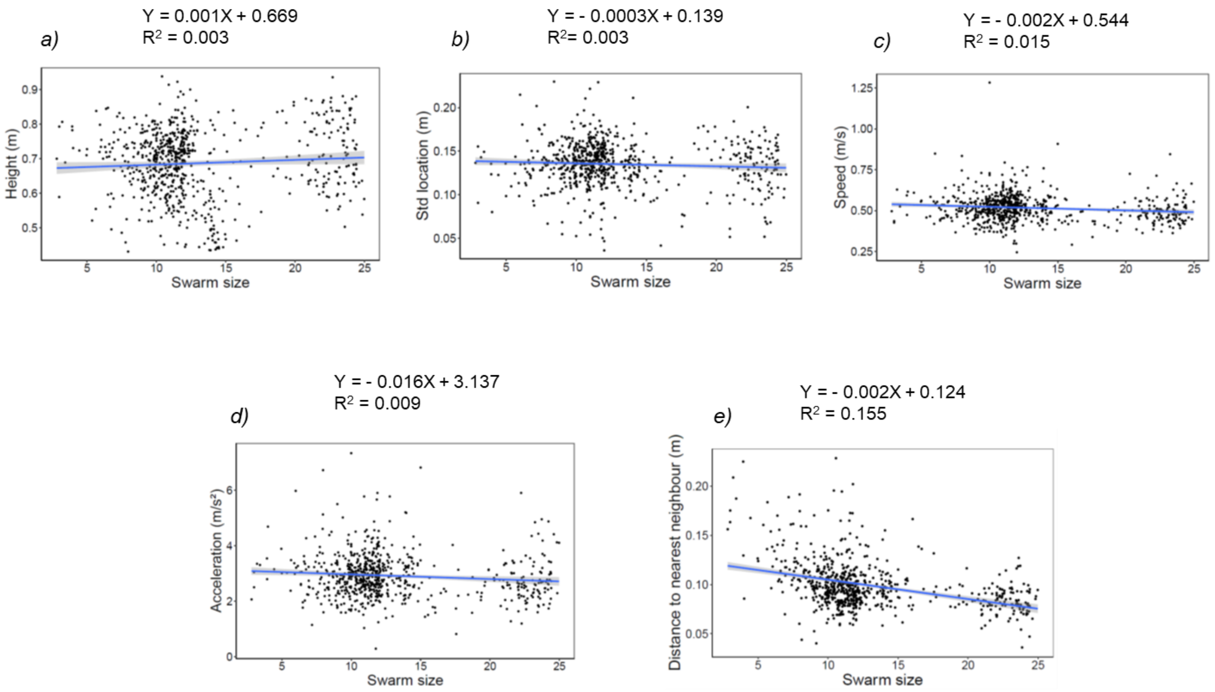


**Figure S6: Effect of the number of swarming mosquitoes (swarm size) on their location and their flight kinematics at the peak phase of the swarming activity.** There was no significant effect of the swarm size on (*a*) the height, (*b*) standard deviation to the mean location, (*c*) flight speed and (*d*) acceleration. However, (*e*) the swarm size significantly affected the distance to the nearest neighbour in the swarm.

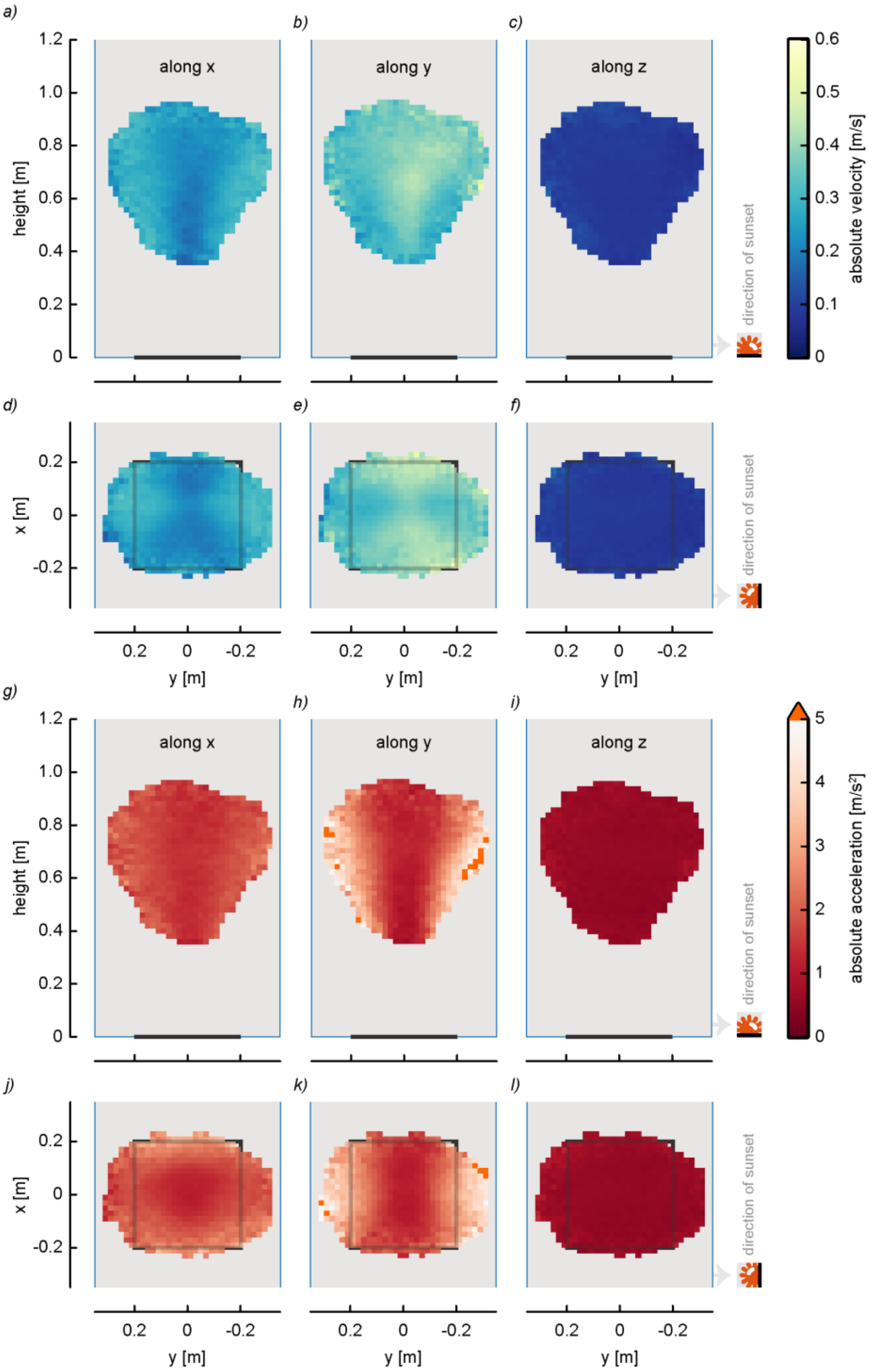


**Figure S7: Spatial distribution of swarming flight kinematics along the three axes.** Side view (first row) and top view (second row) of (*a*-*f*) the spatial distribution of the flight speed and (*g*-*l*) acceleration along the *x*, *y* and *z*-axis. Depending on the viewing plan, the swarm marker is represented by a black line (first row) or a black square (second row). Mosquitoes moved mainly along the *y*-axis (*i*,*e*. in the direction of the artificial sunset horizon) and reached their maximum flight speed in the middle of the swarm.

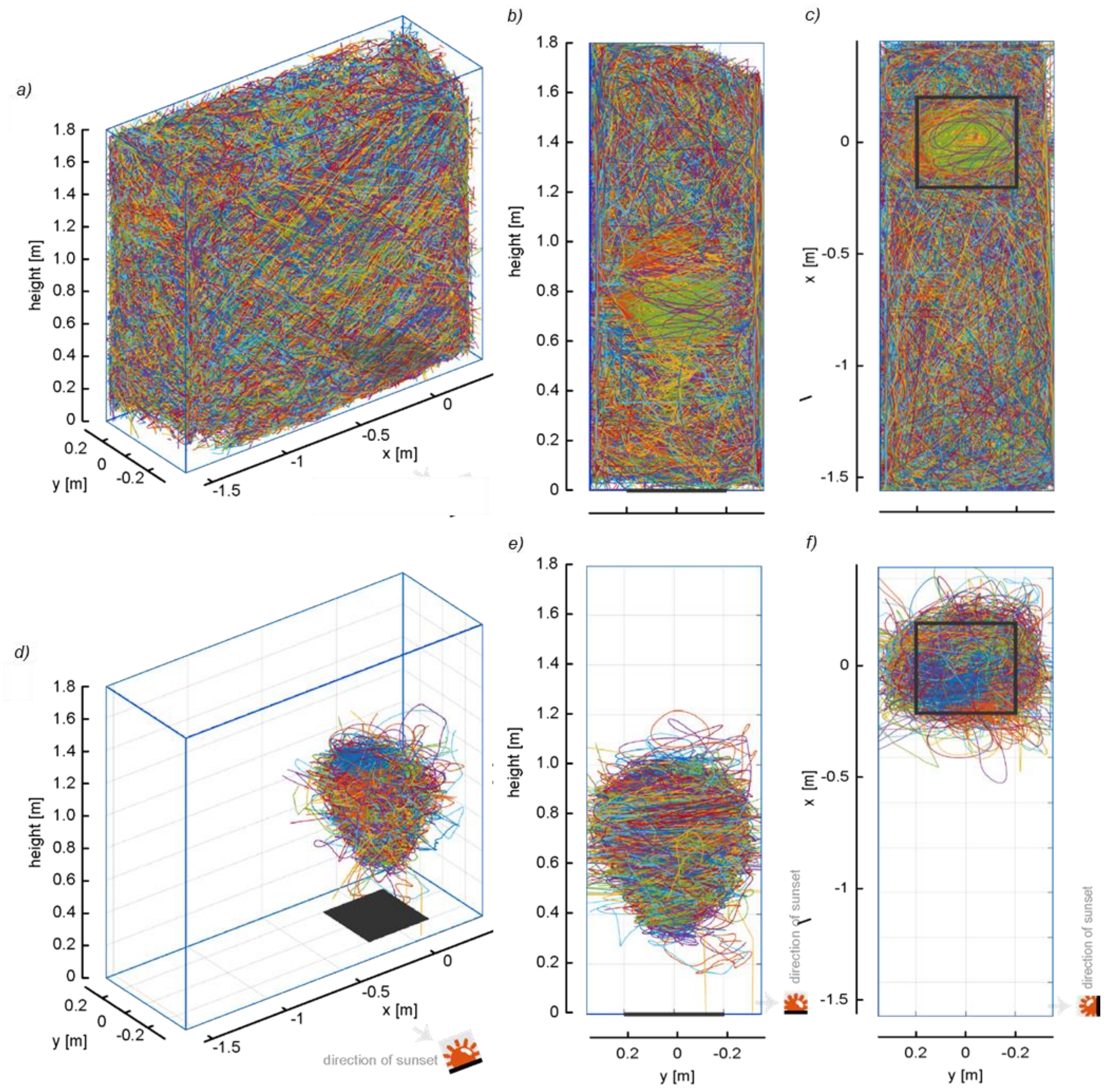


**Figure S8: Raw and filtered three-dimensional tracks.** (*a*) Three-dimensional view, (*b*) side view and (*c*) top view of all the tracks recorded during the six swarming events. (*d*) Three-dimensional view, (*e*) side view and (*f*) top view of filtered swarming tracks. Each colour corresponds to a track. As there was a large number of tracks, colours are used for multiple tracks. Depending on the viewing plan, the swarm marker is represented by (*a*,*d*) a filled-in black square, (*c*,*f*) an empty black square or (*b*,*e*) a black line. Mosquitoes flew throughout the experimental flight arena and the swarming flight activity was located above the marker.

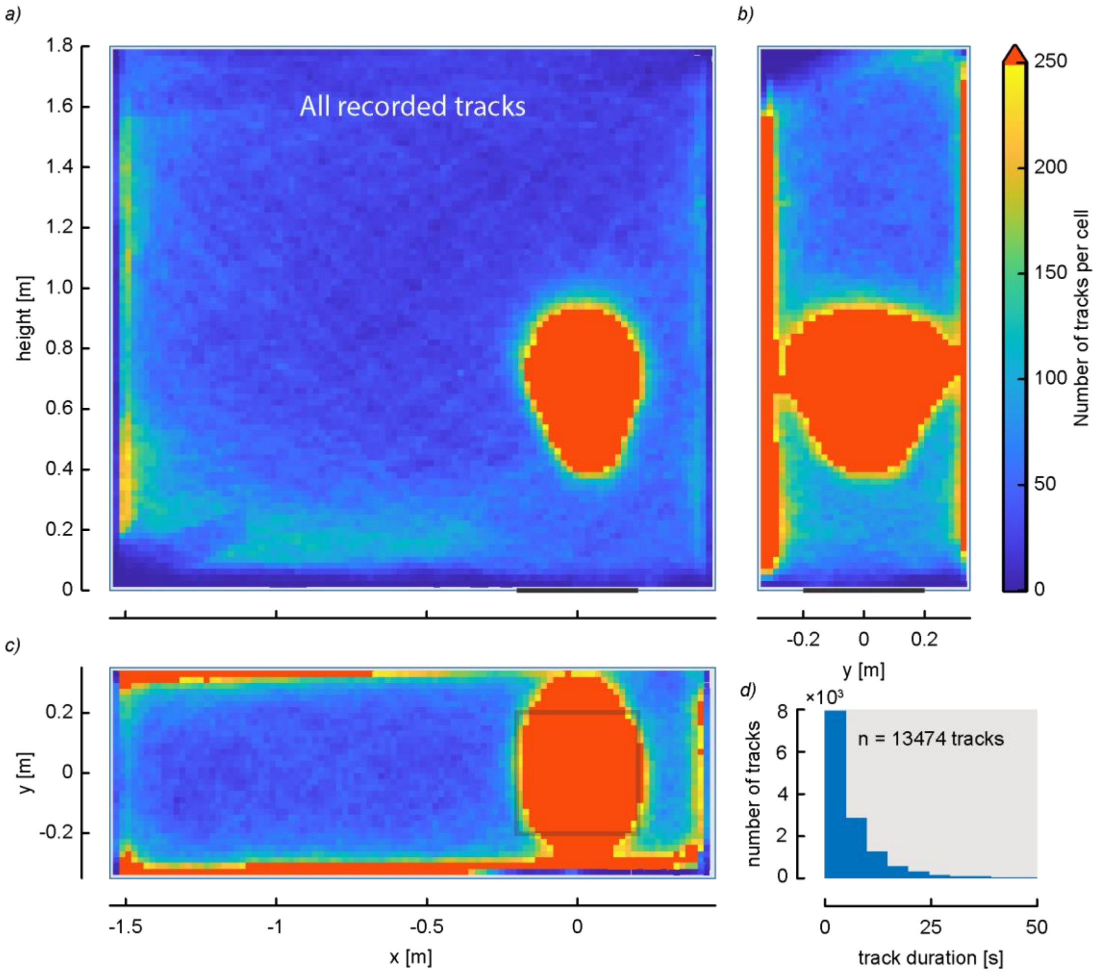


**Figure S9: Spatial distribution of track density for all the recorded tracks.** (*a*) Front view, (*b*) side view and (*c*) top view of the spatial distribution of the number of all the tracks recorded in the flight arena during the swarming periods. Depending on the viewing plan, the swarm marker is represented by (*c*) a black square or (*a*,*b*) a black line. The main flight activity was concentrated over the marker and along the walls of the box corresponding to swarming mosquitoes and to those trying to escape from the box at the ending period, respectively. (*d*) Histogram of the track durations of all the flight activity. Track durations ranged mainly between 0 and 5 seconds.

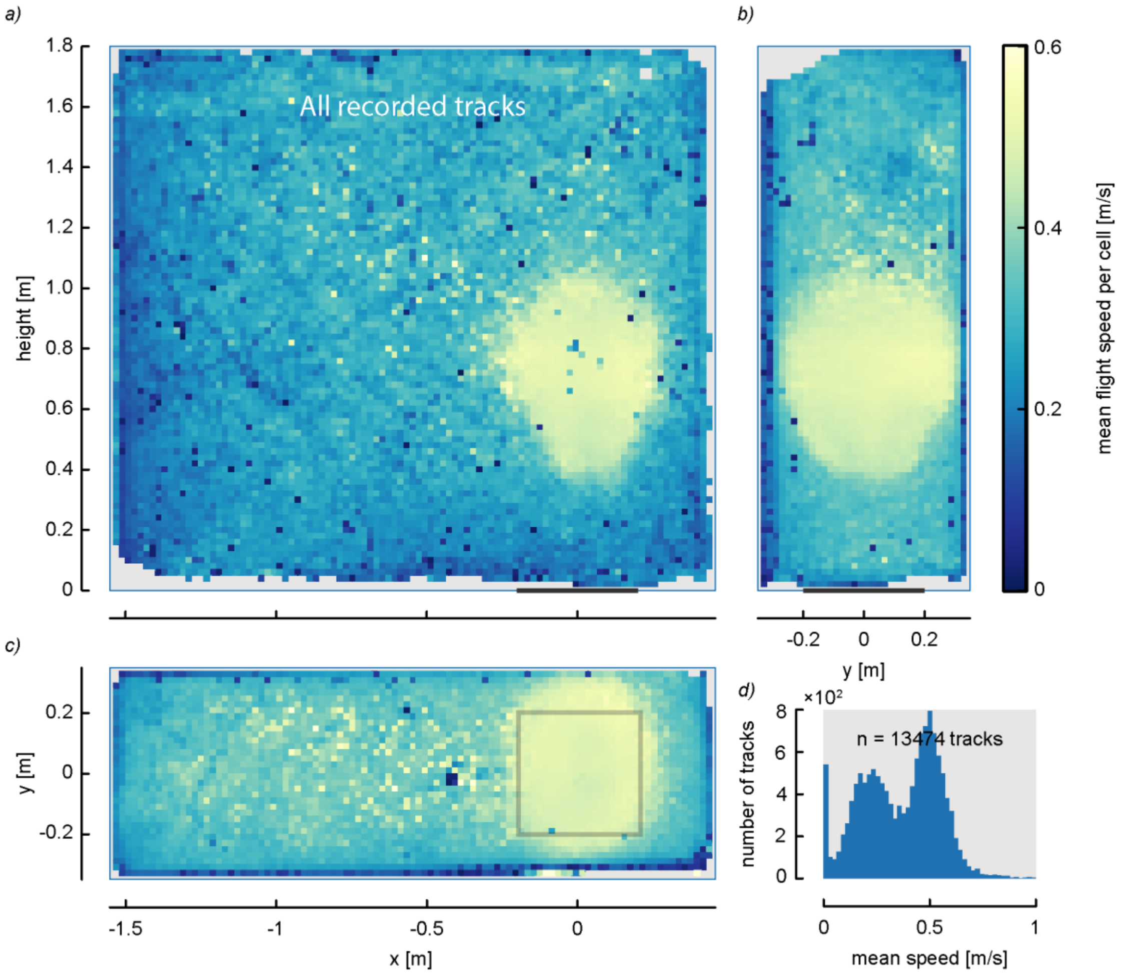


**Figure S10: Spatial distribution of flight speed during the all flight activity.** (*a*) Front view, (*b*) side view and (*c*) top view of the spatial distribution of mosquito flight speed. Depending on the viewing plan, the swarm marker is represented by (*c*) a black square or (*a*,*b*) a black line. The speed at the swarm location was higher than outside the swarm volume. (*d*) Histogram of the mean flight speed per track shows two main flying behaviours: one with a mean flight speed close to 0.25 m/s, likely the random flying mosquitoes, and one with a mean flight speed close to 0.5 m/s, likely the swarming mosquitoes.

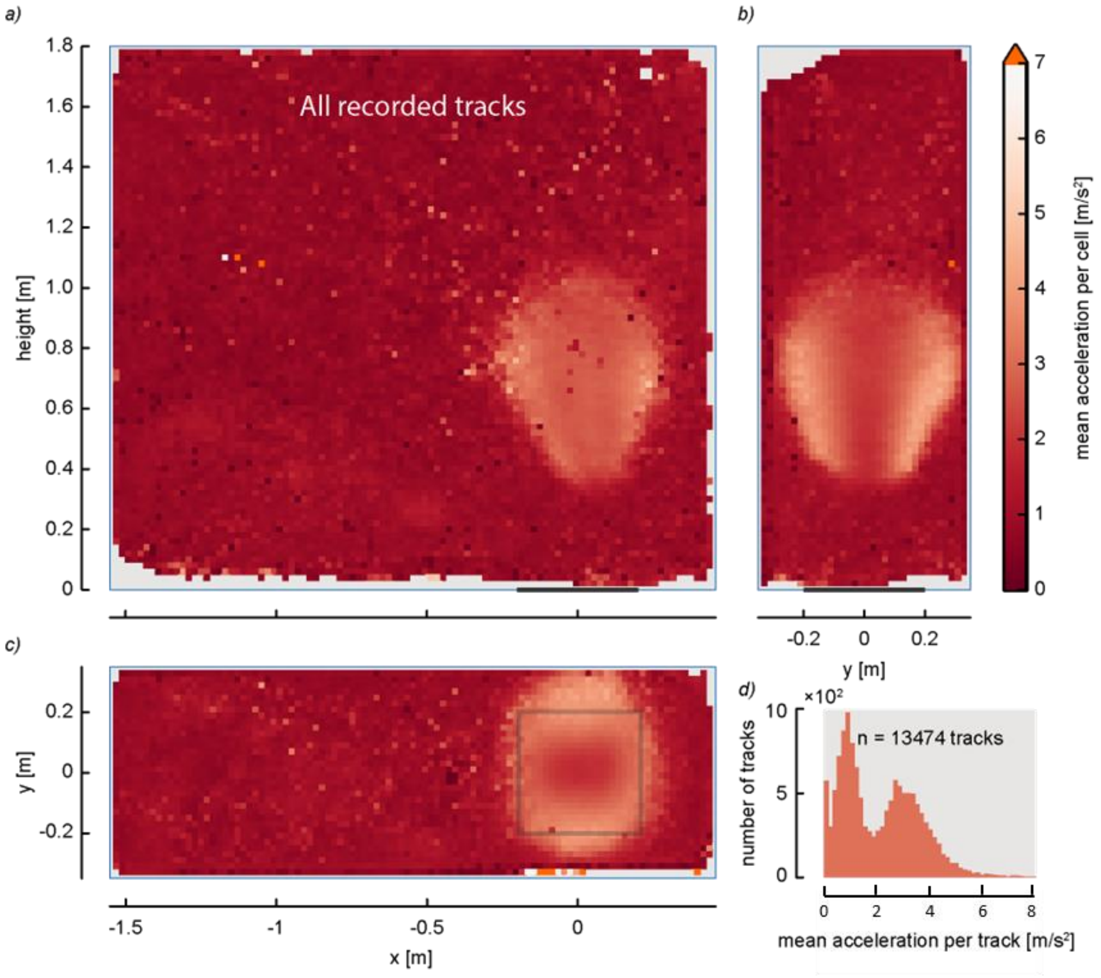


**Figure S11: Spatial distribution of flight acceleration during the all flight activity.** (*a*) Front view, (*b*) side view and (*c*) top view of the spatial distribution of mosquito flight acceleration. Depending on the viewing plan, the swarm marker is represented by (*c*) a black square or (*a,b*) a black line. The accelerations in the swarm volume were higher than outside the swarm. (*d*) Histogram of the mean acceleration per track shows two main behaviours: one with a mean acceleration close to 1 m/s$^2$, likely the random flying mosquitoes, and one with a mean acceleration close to 3 m/s$^2$, likely the swarming mosquitoes.

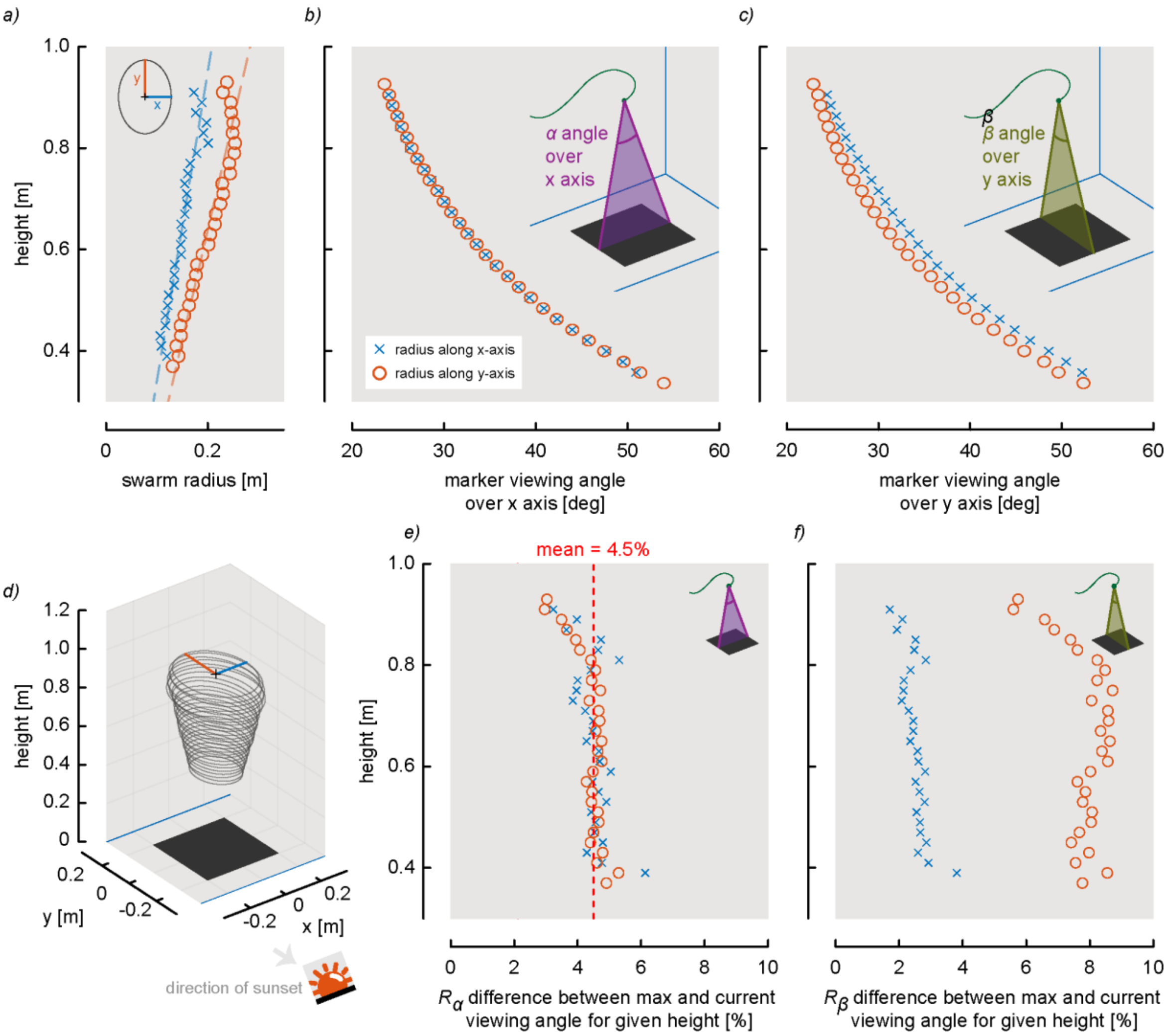


**Figure S12: How is the marker viewed at the contour of the swarm?** (*a*) The swarm contour can be represented as ellipses with varying radiuses over the height. Here, the *x* and *y* radiuses are equal to half the swarm depth for a given height. (*d*) The *x* and *y* radiuses are increasing linearly with the height. (*b*,*c*) Marker viewing angles *α* along *x*-axis or *β* along *y*-axis are computed for the previously mentioned radiuses over various heights. We notice that for a given height, the marker viewing angles *α* over the *x*-axis are the same for both *x* and *y* radiuses (*i.e.* the blue crosses overlay the red circles on panel b). This suggests that the viewing angle over the *x*-axis is consistent along an *xy*-plane ellipse for a given height. This is not true for the viewing angle *β* over the *y*-axis (*c*). (*e*,*f*) The percentage differences $R_\alpha$ and $R_\beta$ between the maximum viewing angle (along *x* or *y*-axis) and viewing angle at a given height ($R_\alpha = (\alpha_{max}-\alpha)/\alpha_{max}\times 100\%$). $R_\alpha$ is constant for the viewing angle over *x*-axis and equal to 4.5%. Using these findings, a proposed model predicting a three-dimensional location and shape of a swarm as a function of 40 cm × 40 cm marker viewing angles, fits well the mean shape of the recorded swarms.
